# Phantom oscillations in principal component analysis

**DOI:** 10.1101/2023.06.20.545619

**Authors:** Maxwell Shinn

## Abstract

Principal component analysis (PCA) is a dimensionality reduction technique that is known for being simple and easy to interpret. Principal components are often interpreted as low-dimensional patterns in high-dimensional data. However, this simple interpretation of PCA relies on several unstated assumptions that are difficult to satisfy. When these assumptions are violated, non-oscillatory data may have oscillatory principal components. Here, we show that two common properties of data violate these assumptions and cause oscillatory principal components: smoothness, and shifts in time or space. These two properties implicate almost all neuroscience data. We show how the oscillations that they produce, which we call “phantom oscillations”, impact data analysis. We also show that traditional cross-validation does not detect phantom oscillations, so we suggest procedures that do. Our findings are supported by a collection of mathematical proofs. Collectively, our work demonstrates that patterns which emerge from high-dimensional data analysis may not faithfully represent the underlying data.

## Introduction

In the age of big data, high-dimensional statistics have risen from a novelty to a necessity. Dimensionality reduction methods aim to summarize high-dimensional data in just a few dimensions, and expose simple low-dimensional patterns embedded within high-dimensional space. One of the simplest dimensionality reduction methods is principal component analysis (PCA) [31, 49]. PCA is useful because it can be computed quickly using a compact matrix equation, and can be understood without extensive mathematical training. It has gained popularity in neuroscience because the output can be interpreted as low-dimensional patterns embedded within high-dimensional data, such as trajectories through time, or spatial patterns. Interpreting PCA in this way has led to numerous neuroscientific discoveries about diverse phenomena [12, 25, 35, 42, 44, 55, 58–60, 69].

While many types of low-dimensional patterns are common in neuroscience, one of the most salient patterns is oscillations. Oscillations in the brain have been studied for over a century [13] and play a crucial role in nearly every brain system in which they can be measured, occurring in both time and space [9, 28, 62]. While oscillations are fundamental to understanding brain function, prior work has warned about a complicated relationship between oscillations and PCA [4, 40, 46, 51]. Strikingly, these studies reported examples of non-oscillatory data that had oscillatory principal components. If this phenomenon is widespread across a variety of data, it could substantially change the interpretation of PCA. In order to trust the results of PCA, it is urgent to understand what causes these oscillations.

Here, we show how and why PCA yields oscillations when no oscillations are present in the data. These “phantom oscillations” are a statistical phenomenon that explains a large fraction of variance despite having little to no relationship with the underlying data. We find two distinct properties of data which cause phantom oscillations through different mechanisms. For each, we provide mathematical proofs and examples demonstrating their impact on data analysis. Finally, we show why traditional cross-validation is ineffective at controlling for phantom oscillations, and propose alternative methods to do so. Collectively, this work illustrates how high-dimensional data analysis is intrinsically linked to low-level properties of the data.

## Results

### Phantom oscillations in PCA

Principal component analysis produces a low-dimensional representation of a high-dimensional dataset (Figure 1a). It takes as its input a set of observations, each consisting of several features, in the form of a matrix. For example, an observation may be a neuron or brain region, a subject, or a trial, and each feature may be the point in time at which that measurements was made. Each “principal component” (PC) consists of (a) a vector of “loadings” or an “eigenvector”, which correspond to patterns in the features; (b) a vector of “scores”, or “weights”, corresponding to patterns in the observations; and (c) the variance explained by the cross product of these two patterns, also known as the “eigenvalue”. PCA is computed by finding the eigenvectors and eigenvalues of the covariance matrix of the data (Figure 1b). The scores are computed by multiplying the data matrix by the loadings, equivalent to a projection of the data onto the eigenvectors (Figure 1c). In what follows, unless otherwise specified, we will focus on datasets of timeseries, where each observation is a neural signal, and each feature is a point in time. In the supplement, we also consider the opposite case of the transposed data matrix, meaning each observation is a point in time and each feature is a neural signal.

**Figure 1:**
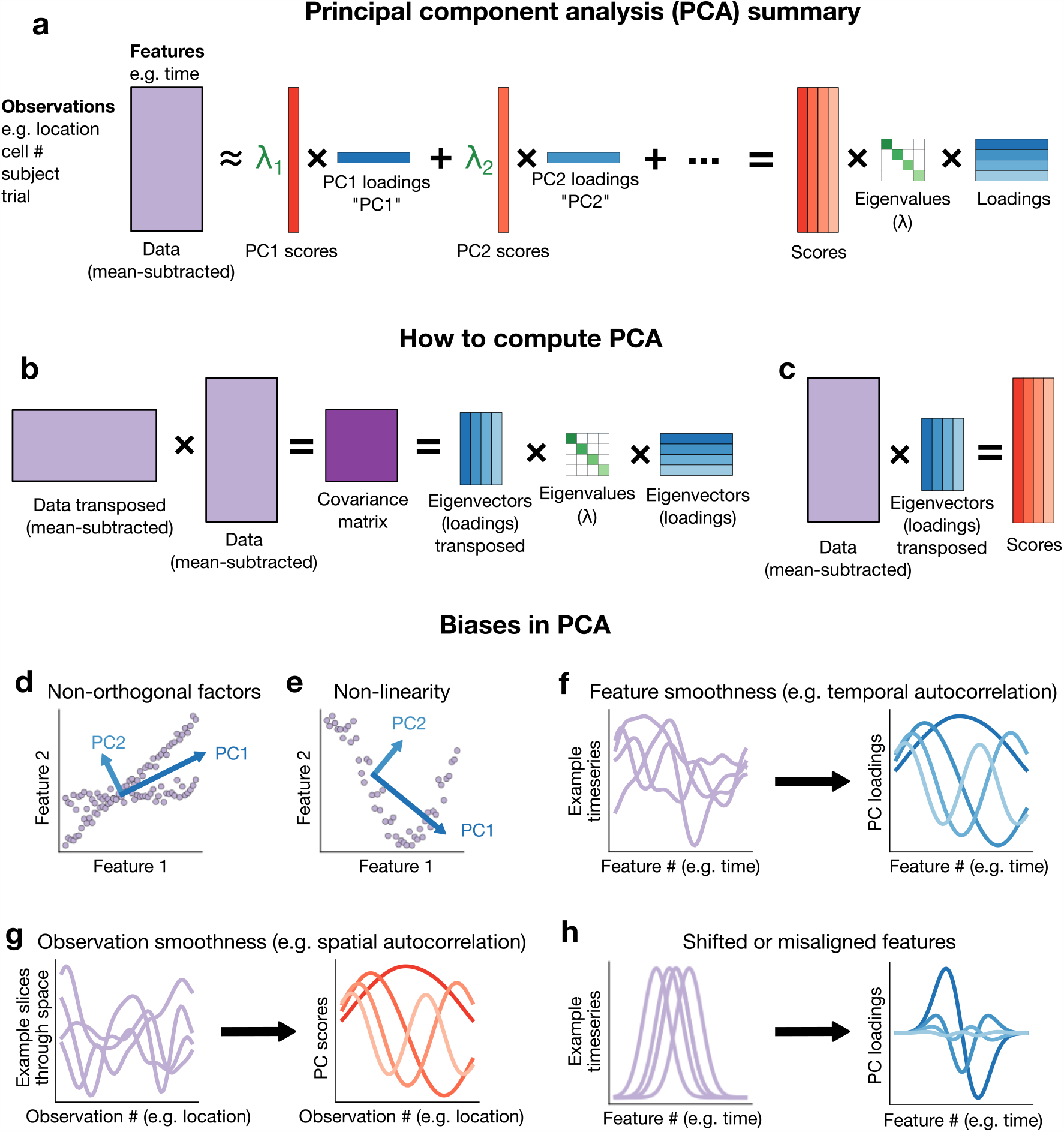
Summary of biases in PCA. (a) Schematic including terminology we use throughout. (b) Summary of how to compute principal components (PCs). (c) Summary of how to compute scores. (d-h) Illustrations of (d) non-orthogonality bias, (e) non-linearity bias, (f) smooth features leading to oscillatory loadings, (g) smooth observations leading to oscillatory scores, and (h) time-shifted features leading to oscillatory loadings.

While PCA itself does not make assumptions about the data, the most popular interpretation of PCA does. This classic interpretation posits that patterns in the loadings or scores indicate some latent underlying patterns in the data. This interpretation is often accurate, but it relies on four assumptions, and may be misleading if those assumptions are violated. First, the underlying patterns must exist and be independent of one another. Second, they must combine linearly to form the observed data. Third, the data must contain exclusively these patterns and additive, uncorrelated noise. Fourth, observations must be independent. When any of these does not hold, the above interpretation of PCA may not hold.

What happens if these assumptions are violated? Violations of the first and second assumptions— independence and linearity of patterns—are known to impact data analysis [18] and are discussed at length in statistics textbooks [34]. Namely, when the latent underlying patterns are not independent of each other, PCA will output components which fail to capture any of the correlated factors (Figure 1d), and if the underlying components are not linear, than PCA will find the best linear approximation (Figure 1e). However, less effort has gone into understanding violations of the final two assumptions: exclusivity, and independence of observations [71].

The third and fourth assumptions are critical to understand because they are violated by most neuroscience data. There are multiple ways these assumptions may be violated. For example, both assumptions are violated by smoothness. Data are smooth across time, or temporally autocorrelated, if nearby points in time have similar values (Figure 1f, left). Smoothness across time is a pattern which cannot be easily expressed as an additive sum, violating the third assumption. As a result, when PCA is performed on smooth timeseries, it will not exhibit interpretable components, because no single pattern can express smoothness across time. Instead, components will exhibit oscillations in the loadings (Figure 1f, right) (mathematical explanation and proofs in Methods). Likewise, smoothness across space, or spatial autocorrelation, occurs when observations are made at spatial locations, and nearby locations are more similar to each other than distant locations. If nearby locations are similar to each other, they will be correlated, which violates the fourth assumption that observations are independent. Smoothness across space produces oscillations in the scores (Figure 1g).

Another common example in neuroscience which violates both the third and fourth assumptions is when observations are similar to each other but shifted in time or space (Figure 1h, left). In other words, there is some underlying pattern in the data, however, this pattern occurs at a slightly different time or position in different observations. This might occur if the timing of some event can vary slightly, or if alignment is imprecise. When this occurs, a linear combinations of features cannot succinctly represent the pattern, which violates the third assumption. In addition, in violation of the fourth assumption, observations are no longer independent, since the magnitude of the random shift in time or space is the primary difference between observations. This type of non-independence will result in loadings that resemble temporally- or spatially-localized sinusoids or wavelets (Figure 1h, right) (mathematical explanation and proof in Methods). Smoothness and shifts are ubiquitous across timeseries and imaging data, and both lead to oscillations in PCA.

We define “phantom oscillations” as oscillations in the PCs of data that vary along a continuum. In most cases, the continuum will be time or space. In one dimension, such as timeseries, phantom oscillations resemble sine waves or localized wavelets, which become Lissajous-like neural trajectories (Figure S1) when plotted against each other. In multiple dimensions, they resemble modes of vibration like a stationary or propagating wave, dependent on the spatial geometry of how they are sampled. Phantom oscillations may also occur on any continuum, such as a graph or a manifold in high-dimensional space (Figure S2). In what follows, we explore two distinct sources of phantom oscillations—smoothness and shifts—and show how they confound neuroscience data analysis.

### Phantom oscillations caused by smoothness in time or space

Phantom oscillations arise naturally in smooth data. Almost all timeseries exhibit some amount of smoothness, so in the absence of true underlying patterns, PCA will yield oscillations. This effect can be observed on both simulated and real data. When we generated synthetic smooth timeseries using low-pass filtered white noise (Figure 2a, top) and then performed PCA, we observed that the loadings resembled sinusoids (Figure 2a, top). The PCs for these smooth timeseries explain much more variance than the PCs for independent unsmoothed white noise timeseries, a common null model for determining the significance of PCs (Figure 2c, top, solid lines). The same results held when different methods were used to generate random smooth timeseries (Figure S3) and when the timeseries matrix was transposed before performing PCA (Figure S4). We repeated this analysis on resting state fMRI data from the Cambridge Centre for Ageing and Neuroscience (Cam-CAN) [64]. To ensure no temporal structure was present across observations, we selected random segments from random brain regions and random subjects, each 118 seconds long (Figure 2a, bottom). We again observed sinusoidal principal components (Figure 2b, bottom) which explained more variance than independent white noise timeseries (Figure 2c, bottom).

**Figure 2:**
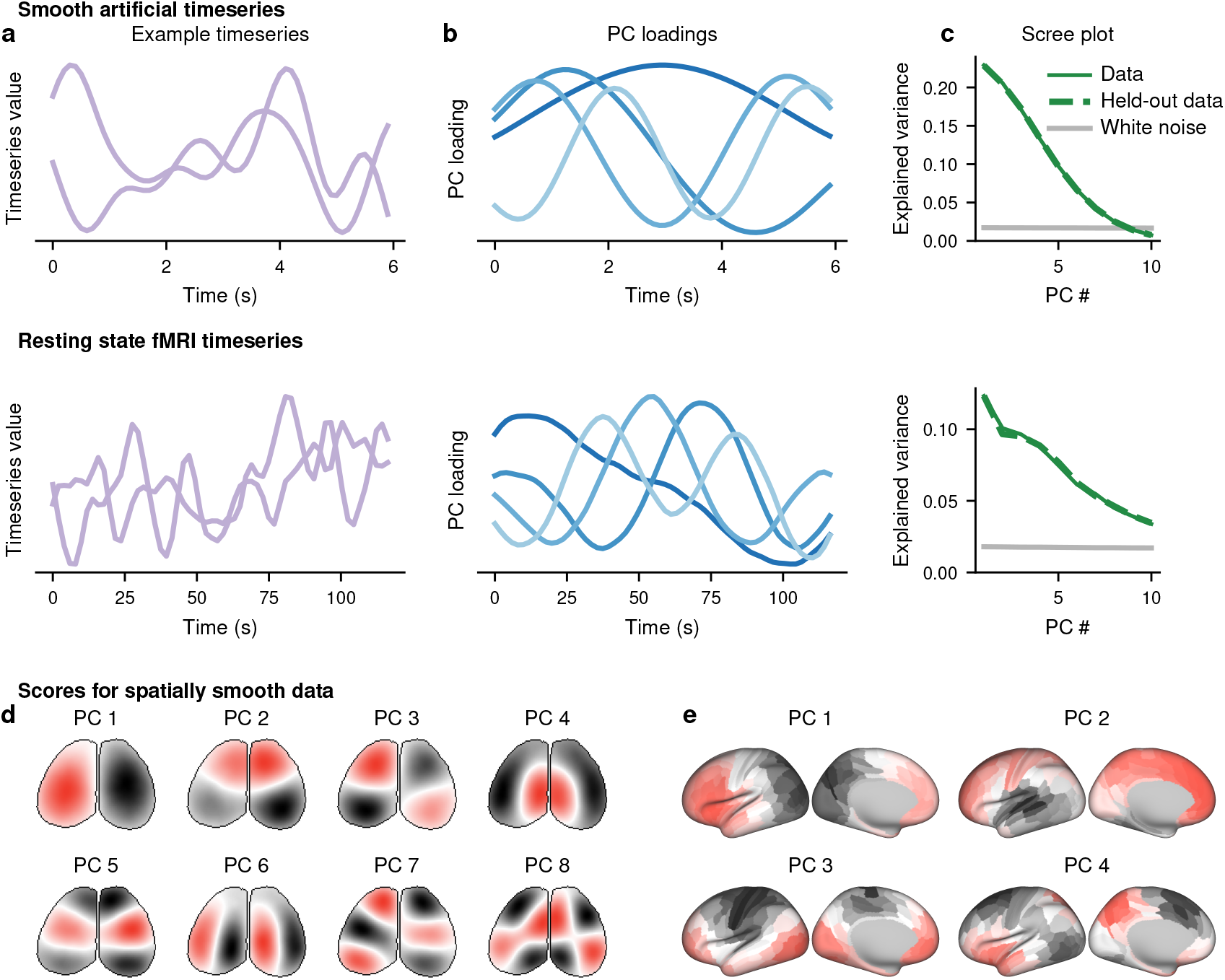
Smoothness causes phantom oscillations in PCA. (a) Example timeseries for smooth artificially-generated data (top) and randomly-selected segments from a resting-state fMRI scan (bottom). (b) Loadings of the first four PCs from large populations of the artificial (top) or fMRI (bottom) data. (c) Variance explained by the first 10 PCs for the artificial (top) or fMRI (bottom) data. Solid green indicates the data from (a) and (b), and dashed green indicates a second independent sample of the size from the same data. Grey indicates the variance explained by the first 10 PCs of an equally-sized dataset of independent white noise. (d-e) Artificially-generated white noise was smoothed across the spatial, but not temporal, dimension in the geometry of a widefield image (d) or a parcellated cortical surface (e). PC scores are plotted on these geometries.

These effects are not the result of overfitting, and do not disappear after traditional cross-validation. To test whether cross-validation could ameliorate this effect, we generated a new independent synthetic dataset of equal size, and sampled new random sequences of the Cam-CAN data. Then, we projected the new data onto the principal components from the original data. We found that the variance explained on the unseen data was approximately equal to the amount explained on the original training set data on both the synthetic and fMRI (Figure 2c) data. In other words, the same oscillatory principal components could explain variance in unseen data, despite the fact that the data were independent by construction. This paradoxical result can be explained by the fact that both halves of each dataset were similarly smooth, so both could be explained by the same set of oscillatory principal components. This indicates that cross-validation cannot control for smoothness-driven phantom oscillations.

In addition to smoothness across time, smoothness across space can drive phantom oscillations. In smoothness across space, nearby locations will be more similar to each other, whether or not there is any smoothness across time. Spatial smoothness is ubiquitous in data such as EEG, widefield imaging, and fMRI. We generated spatially-smooth noise on the dorsal surface of the mouse brain, as in widefield imaging (Figure 2d), and on a single hemisphere of the human cortical surface, as in MRI (Figure 2e). In these simulations, the timeseries were not smooth, but nearby locations in the brain were more similar to each other than distant locations. Since the spatial dimension, not the time dimension, is continuous, we will look for phantom oscillations in the PC scores instead of the loadings. We found that oscillatory-like patterns emerged in the PC scores across the surface of the brain, reminiscent of geometric oscillatory modes [5, 7, 48, 56]. Moreover, some of these patterns were biologically-meaningful, such as localization across medial-lateral and anterior-posterior axes, even though these patterns were not present in the noise. Oscillatory patterns also occur in non-Euclidean topologies, such as a branching latent manifold (Figure S2), due to its relationship with the modes of oscillation (see Methods). Therefore, smoothness in many forms will generate phantom oscillations.

### Phantom oscillations caused by shifts in time or space

A second type of phantom oscillation occurs when data are shifted in time or space. The alignment of a signal in time or space can have a profound influence on PCA. For example, if a fixed underlying pattern occurs in each timeseries, but occurs at a slightly different time in each, then the PCs will display phantom oscillations (Figure 3a, left). These phantom oscillations, shift-driven phantom oscillations, are distinct from those driven by smoothness. Unlike smoothness-driven phantom oscillations, shift-driven phantom oscillations are localized in time around the most prominent peaks and valleys of the signal (Figure 3a, right). These oscillations approximately resemble the rate of change, the first and higher-order derivatives, of the average signal. The phantom oscillations also occur when the matrix was transposed before performing PCA (Figure S5). For most realistic signal shapes, shift-driven phantom oscillations will appear in the form of a spatially localized oscillation.

**Figure 3:**
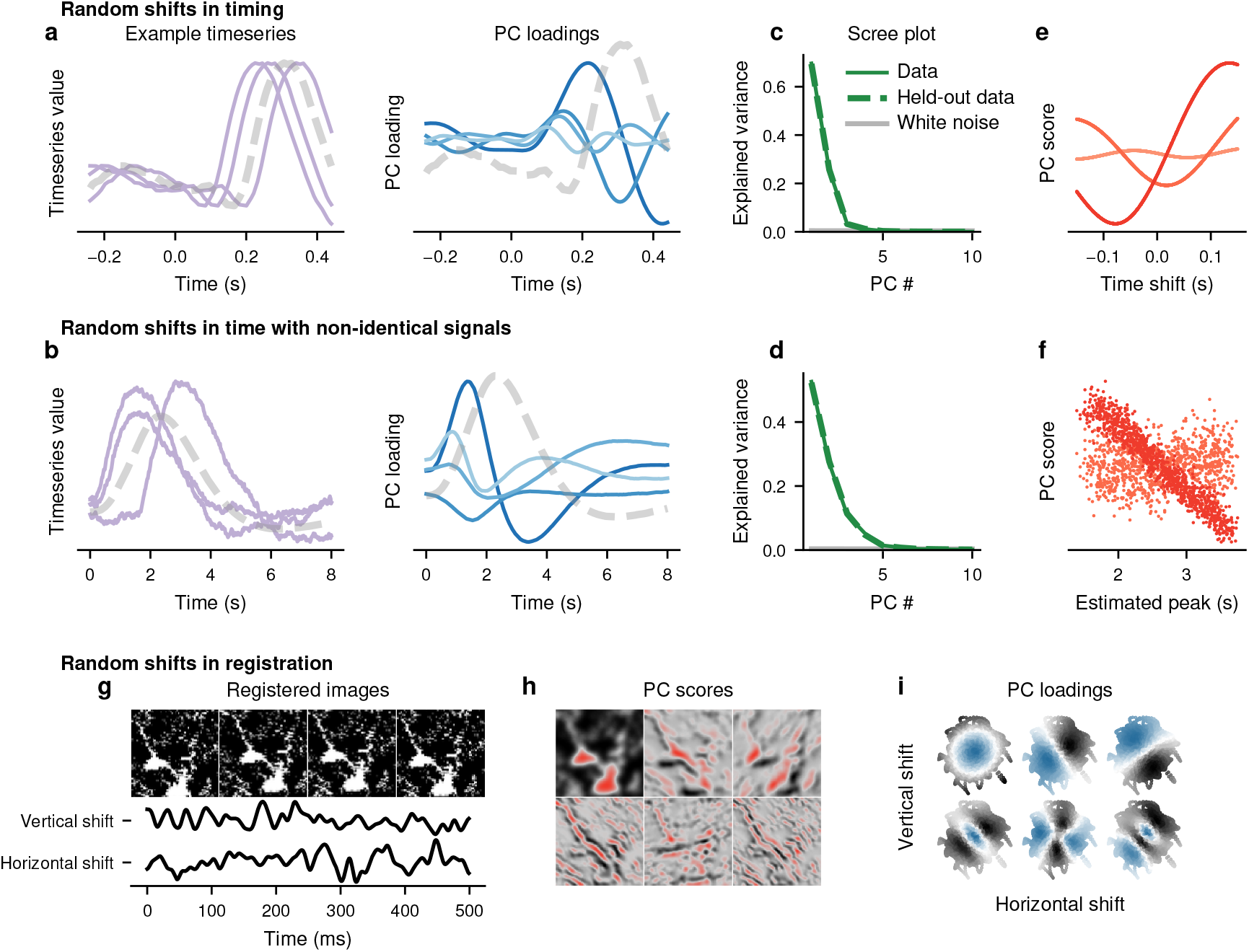
Shifts cause phantom oscillations in PCA. (a) A movement-like signal (dotted grey) was shifted in time according to a centered uniform distribution (light purple). The principal components (PCs) were computed from these artificial data (blue), with the darkest blue indicating PC1 and the lightest PC4. (b) Same as (a), except with a noisy difference of gammas signal which varied parametrically across the gamma parameters and with different noise seeds. (c-d) The scree plot for (a) and (b). (e-f) The shift in time of each signal is plotted on the x axis against the PC score for the data from (a) and (b). (g) Example point spread functions, shifted by up to four pixels in any direction and rescaled by up to 5%. (h) The principal components of the point spread function. (i) An example image of two neurons from two-photon imaging, shifted by up to four pixels in any direction and rotated by up to 4^*°*^. (j) The principal components of the image.

The underlying shifted signals do not need to be identical. We simulated a function inspired by the hemodynamic response function using a difference of gammas with variable time shifts, scaling parameters, and mixing ratios, as well as added noise (Figure 3b, left). We likewise found phantom oscillations in the principal components (Figure 3b, right).

As with smoothness-driven phantom oscillations, traditional cross-validation does not protect against shift-driven phantom oscillations. For the two cases above, we generated a second independent dataset using the same methodology. We projected these new data onto the PCs derived from the original data. We found that the variance explained in the two halves was nearly the same, and was substantially higher than white noise (Figure 3c,d).

One important property of shift-driven phantom oscillations is the relationship of the PC scores with the magnitude of the shift. Since each observation is shifted in time, we can measure the magnitude of the shift relative to some reference. In our simulation where the timeseries are noiseless and where the time shift can be perfectly estimated, the relationship between the PC scores and the time shift will itself be oscillatory (Figure 3e). In more realistic scenarios, the time shift will be estimated noisily for each observation, so the relationship may be more difficult to observe. In our noisy simulation, the relationship appears linear and only present in the first PC (Figure 3f). Nevertheless, there is a clear relationship between the magnitude of the shift and the PC score.

In addition to shifts across time, shifts across space may also drive phantom oscillations. We consider a single frame from a two-photon microscope which we have artificially shifted horizontally and vertically by a few pixels to simulate differences in alignment or registration (Figure 3g). Since the shifts are in the spatial (observation) dimension rather than the time (feature) dimension, we should expect to see phantom oscillations in the PC scores instead of the loadings. Indeed, when we perform PCA on the shifted frame, we find oscillations in the scores spatially localized around the sharpest edges of the frame in both the horizontal and vertical directions (Figure 3h). We already saw that, in the temporal case, the PC scores showed a strong relationship with the magnitude of the shift. Here, there are two dimensions to the shift: horizontal and vertical. For each component, we can examine a scatterplot of the horizontal and vertical shift, where points are colored by the value of the PC loading for those shifts. We see a strong oscillatory pattern relating the PC loadings to the shift (Figure 3i). Therefore, shifts across time or space can lead to phantom oscillations.

### Phantom oscillations in a decision-making experiment

To show how phantom oscillations manifest in real data, we examine a classic random dot motion perceptual decision-making experiment with recordings in monkey lateral intraparietal cortex (LIP) [57]. In this experiment, monkeys were trained to discriminate the apparent direction of motion of dots located centrally in their visual field. Once they reached a decision, they indicated their decision by directing their gaze to one of two peripherally located targets (Figure 4a). While monkeys were performing this task, a neuron in LIP was recorded such that one of the saccade targets was located in the neuron’s receptive field (target T-IN) and the other was opposite (target T-OUT). This allowed allowed neural activity to be compared when the choice was towards T-IN compared to away from T-IN. Neural activity from different parts of this task is expected to contain both smoothness-driven and shift-driven phantom oscillations.

**Figure 4:**
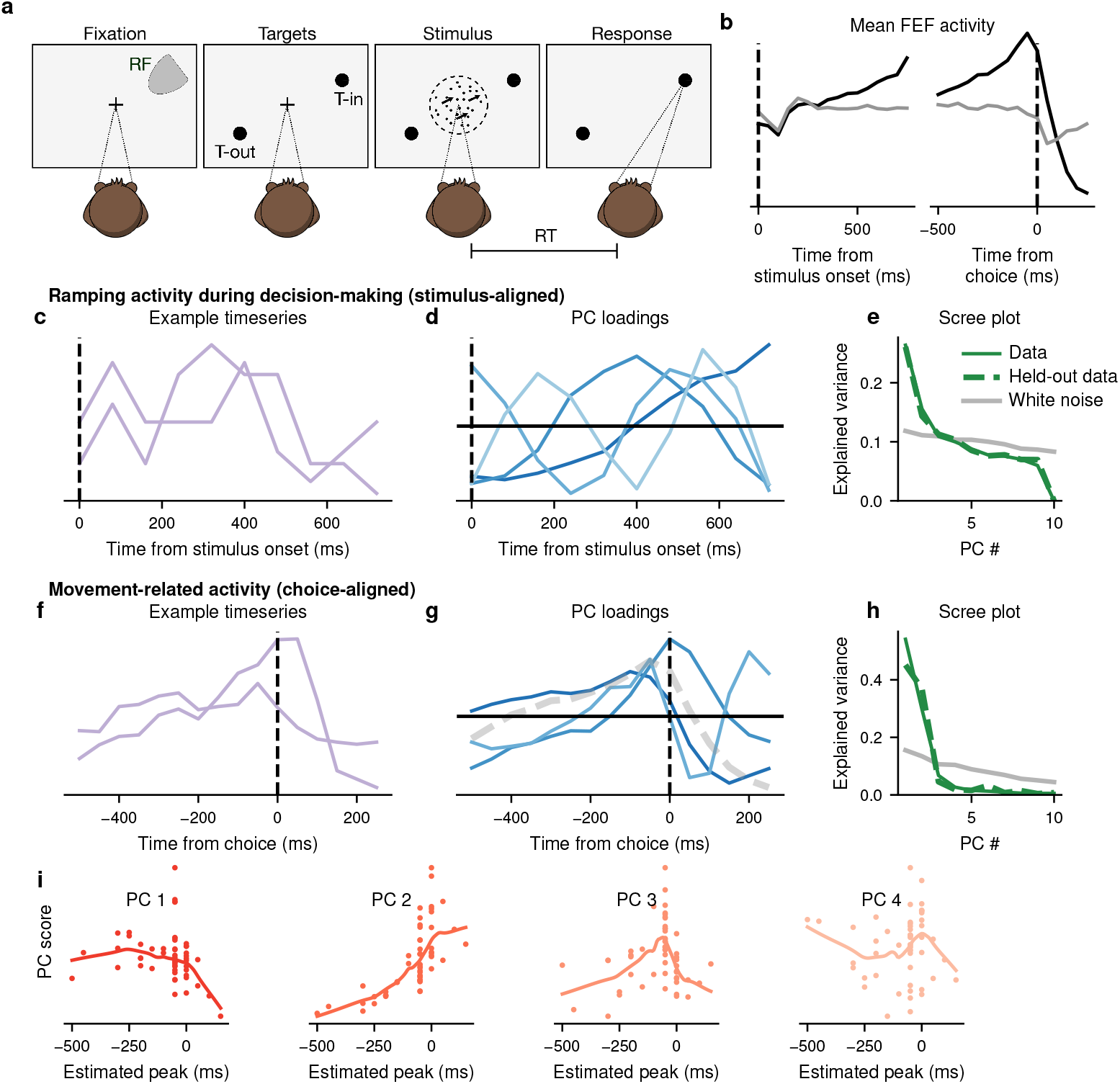
Phantom oscillations in the random dot motion task. Roitman and Shadlen [57] recorded from LIP in monkeys performing a decision-making task. (a) Diagram of the experiment. (b) Mean activity for trials with a long response time (RT) (RT *>* 800 ms) with a choice in the direction of T-IN (black) or T-OUT (grey). (c-e) Timeseries during decision-making period for long trials (RT *>* 800 ms). (c) Two example timeseries during this period, representing individual trials. (d) PCA was performed on all trials and the PC loadings are shown. (e) PCA was recomputed on 1/2 of the trials. Variance explained is shown in solid green. The remaining 1/2 of trials were projected onto the loadings, and the variance explained by this projection is dashed green. Grey is the variance explained by the PCs of an equivalent-sized white noise dataset. (f-i) Timeseries during decision-making period for long trials (RT¿800 ms). (f) Two example timeseries during this period, representing individual cells. (g) PCA was performed on all neurons and the PC loadings are shown. (h) PCA was recomputed on 2/3 of the neurons. Variance explained is shown in solid green. The remaining 1/3 of neurons were projected onto the loadings, and the variance explained by this projection is dashed green. Grey is the variance explained by the PCs of an equivalent-sized white noise dataset. (i) PC scores are plotted against the estimated peak of each neuron’s timeseries. LOESS-smoothed lines indicate the trend.

First, the activity of neurons in LIP leads to smoothness-driven phantom oscillations. Many neurons in LIP are known to reflect the accumulated information in favor of a choice towards T-IN (Figure 4b, left) [57], often modeled using diffusion [53, 66]. Diffusion produces smooth timeseries, and simulations of diffusion show phantom oscillations (Figure S3). Therefore, trial-by-trial neural activity aligned to the onset of the stimulus should show smoothness-driven phantom oscillations. We examine the first 700 ms of activity after the stimulus, using only trials at least 700 ms long (Figure 4c). We find that the principal components of these data closely resemble sinusoidal oscillations (Figure 4d). Using cross-validation, we fit data on a subset of 66% of the data and project the remaining 33% on to these PCs. The variance explained on the held-out data is similar to that of the training data, and much higher than for a white noise timeseries (Figure 4e). Therefore, phantom oscillations are present in neural activity traditionally considered to be diffusion-like.

Second, bursts of activity in response to eye movements lead to shift-driven phantom oscillations. Many neurons in LIP show sharp increases in activity during eye movements towards T-IN (Figure 4b, right). Furthermore, these bursts of activity are unlikely to show exactly the same peak timing in each neuron (Figure 4f). Therefore, the mean activity of cells should show shift-driven phantom oscillations. Indeed, we find that the principal components resemble the derivatives of the mean (Figure 4g). When PCA is performed on 66% of data and the remaining 33% are projected onto the PCs, a similar amount of variance is explained in the training and held-out data, much higher than for a white noise timeseries (Figure 4h). Furthermore, the scores of these first principal components correlate with the peak of the timeseries, an estimate of the shift (Figure 4i). Therefore, both smoothness-driven and shift-driven phantom oscillations are present in different aspects of the same decision-making experiment. This dataset is also relatively small by modern standards, containing only 54 neurons, demonstrating that phantom oscillations are not limited to big data.

### Detecting true oscillations in data

If oscillations are observed in PCA, how can we determine whether they are true oscillations or phantom oscillations? Distinguishing phantom oscillations from true oscillations is difficult, and sometimes, impossible. For example, multiple cycles of a true oscillation must be present in order to distinguish it from a phantom oscillation. Without this, true oscillations are impossible to identify, both on a technical and a conceptual level. In general, PCA is not well-suited to detecting oscillations, so most oscillations observed tend to be phantom oscillations. Nevertheless, we present three strategies to help identify true oscillations in PCA.

To demonstrate these strategies, we analyze an experiment containing repetitive circular movements, designed to induce true oscillations in neural activity. Ames and Churchland [3] instructed monkeys to move through a linear virtual reality environment by moving a manipulandum in a cyclic pedaling motion (Figure 5a). The distance the monkey needed to travel in virtual reality was fixed and corresponded to seven pedals on the manipulandum. While the monkeys performed the task, multiple neurons from primary motor cortex were recorded simultaneously using 24 channel probes. Consistent with Ames and Churchland [3], we discarded first cycle and last two cycles to maintain a steady state cycling motion, rescaling the remaining four cycles to a 2 s interval for analysis (Figure 5b). Individual neurons show oscillatory activity during this task (Figure 5c). The principal components resemble sinusoidal oscillations (Figure 5d) and explain a large fraction of variance, even after cross-validation (Figure 5e). We compare these principal components to those of the phantom oscillations presented previously in simulated data, resting state fMRI, and the random dot motion task. The principal components and scree plots for these data can be viewed in Figure 2, Figure 3, and Figure 4.

**Figure 5:**
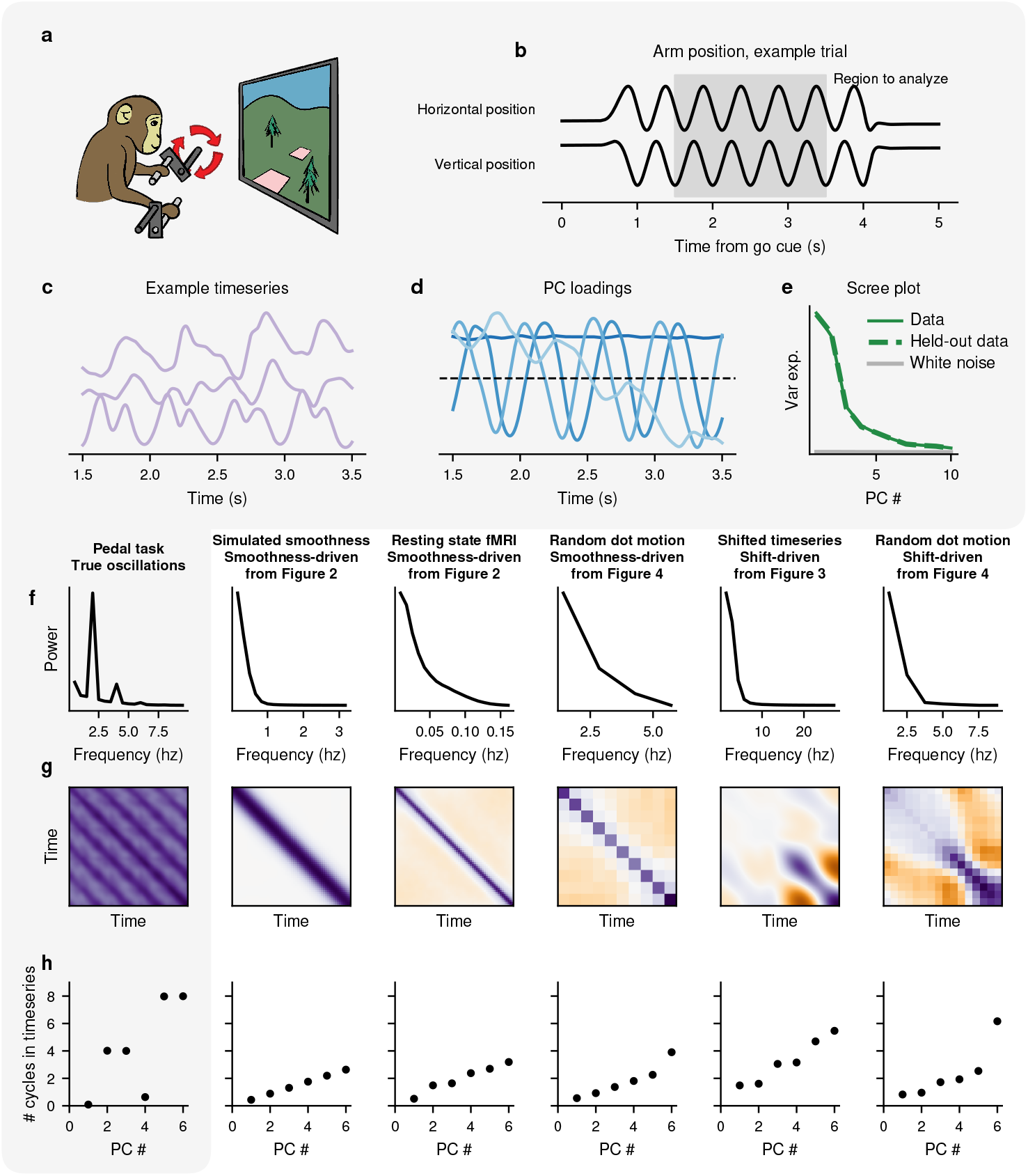
True oscillations in a pedal task. Ames and Churchland [3] recorded from motor cortex in monkeys performing a pedaling task. (a) Diagram of the experiment, reproduced from Ames and Churchland [3]. (b) Horizontal and vertical position of the arm during the pedaling task during an example trial. Only the middle four cycles were analyzed. Cycles were rescaled in this period to allow averaging across trials by arm position. (c) Mean activity of three example neurons during the analyzed time period. (d) PCA was performed on the mean activity of each neuron during the task, and the loadings are shown. (e) PCA was recomputed on half of the neurons, and the variance explained is shown in green (solid). The remaining neurons were projected onto the PCs from the first half, and the variance explained is shown in green (dashed). For comparison, PCA was performed on an equal number of white noise timeseries of the same length, and variance explained is shown in grey. (f-g) Measures for detecting phantom oscillations are shown for the pedal task, a dataset with true oscillations (left) as well as the other datatsets exhibiting smoothness-driven and shift-driven phantom oscillations we have seen so far. (f) The power spectrum from the pedal task. (g) The covariance matrices for each of these datasets. (h) For each dataset, the number of oscillations (i.e., the wave number) for each PC loading is shown.

One strategy to identify true oscillations is to use traditional methods for detecting oscillations in timeseries. True oscillations will be visible using traditional methods, such as peaks in the power spectrum, whereas phantom oscillations will not. In our cycling dataset, we see a clear peak in the power spectrum at 2 hz corresponding to the four oscillations of the manipulandum, as well as a peak at 4 hz corresponding to the first harmonic (Figure 5f, left). By contrast, no such peaks are present in the power spectrum for the other datasets (Figure 5f).

A second strategy to identify true oscillations is to inspect the covariance matrix. In phantom oscillations, we expect one thick prominent stripe along the diagonal of the matrix, covering either all (Figure S3) or part (Figure S6) of the diagonal. By contrast, for true oscillations, we expect multiple diagonal stripes throughout the covariance matrix. These multiple stripes represent the correlation across different phases of the same oscillation. We observe multiple stripes in the pedaling dataset (Figure 5g, left), but only one thick stripe in the others (Figure 5g).

A third strategy is based on the fact that a single phantom oscillation principal component is never found alone. Phantom oscillations always appear across many frequencies, with a distinct frequency in each principal component. Smoothness-driven phantom oscillations with lowest oscillatory frequency will explain more variance than those with higher oscillatory frequency, and shift-driven phantom oscillations tend to follow a similar pattern as well. The slope of this relationship should increase consistently with a slope less than 1, linearly or step-wise, and should not make any large jumps. Therefore, the oscillation frequency will form a monotonic relationship with the ordering of the PCs. In the simulated, fMRI, and decision-making data, we see a near-linear relationship between the PC number and the frequency of the oscillation (Figure 5h). By contrast, in the cycling data, after a first constant PC, the second and third PCs oscillate four times, and the fifth and sixth oscillate 8 times (Figure 5h, left).

Prospective shift-driven phantom oscillations have an additional property which distinguishes them from true oscillations. As we saw previously (Figure 3e,f) (Figure 4i), the PC scores of shift-driven phantom oscillations are correlated with the magnitude of the shift. Computing this correlation requires estimating a time shift from each observation, using methods including cross-correlation, peak detection, or other techniques. Since the relationship may be non-monotonic, a Pearson correlation may be insufficient, but examining this relationship by eye should provide an indication of non-linear relationships. Therefore, several signatures can be used to determine whether an oscillation is a phantom oscillation or a true latent oscillatory signal in the data.

## Discussion

Despite being one of the simplest forms of dimensionality reduction, PCA’s interpretation can be complicated. Here, we showed that PCs may exhibit oscillations that do not exist in the underlying data. This is because real-life data often vary along a continuum—such as time in a timeseries, or position in spatially-embedded data. However, continua often violate the implicit assumptions underlying the classic interpretation of PCA. Other examples of continua include neurons along a linear probe, subjects in a longitudinal experiment, or brain regions in an imaging experiment. We showed that there are two distinct ways that phantom oscillations can arise from a continuum: smoothness, and shifts across observations. Our simulations and data analysis showed different patterns for smoothness-driven and shift-driven phantom oscillations. Likewise, in the Methods, we derived that smoothness is related to the second derivative of the timeseries, whereas shifts are related to the first derivative. We also provided a simple method to test for both types of phantom oscillations. Our results may apply more broadly to topographic space, such as place cells [21], retinotopy and tonotopy, or concepts in high-dimensional space defined across a graph.

We are not the first to observe phantom oscillations. Previous work found similar oscillations from PCA in neural data [51] and tuning curves [75], as well as other types of smooth data, such as natural images [47], music [15], and spatial population genetics [46]. Phantom oscillations also resemble the oscillation modes in whole brain imaging experiments [5, 7, 48, 56], which is connected to PCA through the same set of differential equations (see Methods). Phantom oscillations are also closely related to the “horseshoe effect” found in k-means clustering [38] and multi-dimensional scaling (MDS) [20, 37]. Nevertheless, in the Methods, we show that phantom oscillations are more complicated than simply the terms of a Fourier series. This is because phantom oscillations often have non-integer and non-uniformly-spaced oscillatory frequencies. Additionally, phantom oscillations may only produce one sinusoid at each frequency, whereas a Fourier series always has a sine and a cosine at each frequency. The only condition in which phantom oscillations do resemble a Fourier series is the rare situation when the edges of the timeseries wrap around, such that the first and last points are correlated with each other in the same way as neighboring points. This results in a “step-like” scree plot (see Methods and Figure S3).

It remains to be shown whether it is possible to compute PCA in a way which is robust to phantom oscillations. One potential approach for smoothness-driven phantom oscillations is to formulate an appropriate noise model. For instance, in factor analysis and probabilistic PCA (PPCA), there is an explicit model of the noise [16] which assumes noise is independent and normally distributed [70]. Using a correlated noise model to account for smoothness may help reduce the effect of smoothness-driven phantom oscillations.

The conditions that cause phantom oscillations are ubiquitous in neuroscience data. Smoothness may occur for biological or methodological reasons. On the biological side, reasons include slow intrinsic neural and circuit time constants, and the fact that behavior is often history-dependent. On the methodological side, reasons include measurements of activity with slow dynamics, such as calcium indicators or the hemodynamic response function in fMRI. More generally, any type of low-pass filtering makes data vulnerable to phantom oscillations. Shifts may also occur for biological or methodological reasons. On the biological side, different subjects or neurons may have slight intrinsic differences in response time. Response time may also vary across trials within a subject, or may depend on external factors like attention or arousal [41]. There also may be multiple points of interest, such as the stimulus onset and the movement onset in our decision-making dataset, making it difficult or impossible to align to both. On the methodological side, a large sampling rate or imprecision in alignment may create the same effect. Additionally, volumetric imaging technologies such as two-photon imaging or fMRI usually acquire volumetric planes sequentially, leading to shifts in timing across planes.

PCA is often used as a way to compare brain activity across different experimental conditions. However, if differences in PCs are observed when phantom oscillations are present, these differences may be a proxy for some simpler phenomenon. For example, since the PC score is correlated with the location of the peak in shift-driven phantom oscillations, any change in the peak latency across conditions will be reflected in the PCs. Moreover, anything which is correlated with this latency will also be correlated with the PC score. In smoothness-driven phantom oscillations, differences in smoothness across observations will also be reflected in the PCs. Such differences may arise from many sources, including variation in firing rate or signal-to-noise ratio. Therefore, in the presence of phantom oscillations, group differences in PC loadings or scores can often be explained by a simpler statistical property of the data.

In this work, we focused on the case where time is the feature dimension, and thus, the loadings represent points in time. By contrast, in neuroscience, it is common to perform PCA with time in the observation dimension, which is equivalent to performing PCA of the transpose of the data matrix. We focus on the former for three reasons. First, performing PCA along the feature dimension allows us to assume our observations are independent samples from the same statistical process. The standard interpretation of PCA, where components represent latent patterns in the data, assumes that observations are independent. If smoothness occurred in the observation dimension, then the observations would by definition no longer be independent. Our mathematical analyses are largely based on the Karhunen-Loève transform, which also assumes observations are independent. Second, the relationship between the covariance matrix and smoothness is only visually evident as a thick diagonal when smoothness occurs in the feature dimension. It may be possible to rearrange the rows and columns of the covariance matrix to emphasize this thick diagonal, but this is beyond the scope of most analyses. Third, PCA along the observation dimension gives very similar results to PCA across the feature dimension. We showed that, in our simulations, both smoothness and time shifts cause oscillations regardless of whether the smoothness or time shifts are in the feature or observation dimension. We also provided a mathematical justification for why this is the case in the Methods. Therefore, phantom oscillations occur no matter the dimension that PCA is performed.

PCA is closely related many other forms of dimensionality reduction. The results presented here apply to any method which involves finding the eigenvectors of a matrix multiplied by its transpose, such as a covariance matrix, correlation matrix, or matrix of second moments. For example, our mathematical analyses can be applied almost identically to SVD, which is the same method as PCA except without mean subtraction. Additionally, similar oscillatory biases have been reported for other forms of dimensionality reduction, including demixed PCA [39], jPCA [40], canonical correlation analysis (CCA) [30, 51], partial least squares (PLS) [30], and network analysis of a correlation matrix [65]. Many other studies explore how low-level statistics influence dimensionality reduction [8, 11, 18, 22, 29, 30, 40, 43, 50, 51, 63, 65]. These studies mirror our own by showing that patterns in high-dimensional data analysis may not always reflect true patterns in the data.

## Acknowledgments

We thank Katarzyna Jurewicz and Maxime Beau for feedback on the manuscript, and the members of the Johns Hopkins decision-making group for helpful discussions. This study is funded by the European Molecular Biology Organization grant ALTF 712-2021.

## Materials and Methods

### 1 Mathematical framework

In order to understand the mathematics describing smoothness-driven or shift-driven phantom oscillations in PCA, we must first explore some properties of the tools we will use.

#### 1.1 Fundamental assumptions

We will assume that one dimension of the data varies along a continuum. By this, we mean that the points along this dimension can be thought of as a discretization of some continuous function *f* (*x*). Frequently, this dimension is time, and we will refer to this dimension as “time” for convenience. However, the same results hold if we instead consider other one-dimensional continua in space, such as positions along a linear probe or brain regions along the anterior-posterior axis. The results also hold for smoothness across other measured variables, such as age, or abstract relationships, such as position on a manifold. While our arguments conceptually apply to multi-dimensional continua as well, such as position in x-y space, we mostly focus on the unidimensional case due to its analytical tractability.

Additionally, there must be another dimension to represent observations. This dimension need not vary along a continuum, and may represent subjects, neurons, or other entities. For simplicity, we will assume that each observation is independent. However, in practice, similar results still hold if observations are not independent or if they vary along a different continuum. For example, observations may be trials or sessions with serial correlations, or they may be voxels that are spatially autocorrelated.

We will organize our data into a *m × n* matrix *D*, with *m* observations and *n* timepoints. In other words, the first dimension represents the observations and the second is the continuous dimension. This convention may differ from other treatments.

#### 1.2 Covariance matrices and covariance kernels

Our analysis focuses on covariance matrices. As shown in Figure 1, the PC loadings of any dataset can be found using exclusively the covariance matrix. As a result, two datasets will have the same PC loadings if they share the same covariance matrix, even if the datasets are very different from each other. This relationship also applies in the opposite direction, so if two datasets have the same PC loadings, they must have the same covariance matrix. Therefore, covariance matrices are a convenient way to reason about the relationship between a dataset and its PC loadings.

Another convenient feature of covariance matrices is they can be generalized to continuous time processes through a covariance kernel. A *n×n* covariance matrix *C* represents *n* discrete timepoints, where the element *C*[*i, j*] at the *i*’th row and the *j*’th column signify the covariance between these two timepoints. Another way to think about this is as a function, which takes two timepoints *t*_1_ and *t*_2_ as its input, and returns the covariance at those two timepoints. This function *C*(*t*_1_, *t*_2_) provides all of the information that is provided by the covariance matrix. However, when we think about this as a function, it is no longer necessary for the timepoints *t*_1_ and *t*_2_ to be integers. Instead, we could define this function to accept any values of *t*_1_ and *t*_2_ that fall within the interval [0, *T*]. This function *C*(*t*_1_, *t*_2_), also known as a covariance kernel, indicates the covariance between any two points in time, where *t*_1_ and *t*_2_ fall within the closed interval [0, *T*]. Therefore, we can think of a covariance matrix as a discretization of some underlying covariance kernel. This construction, while impractical for data analysis, will be very useful in the proofs we discuss below.

We assume that all of our observations are independent samples from some underlying statistical process. The covariance matrix or covariance kernel can be used to determine several properties about the statistical processes. If the timeseries is smooth, the covariance matrix will show a thick diagonal stripe, with large values the diagonal and small values farther away from the diagonal. If the process is circular, i.e., the edges wrap around, then the values at the opposite corners of the matrix, as far away as possible from the diagonal, will be positive. If the process is isotropic, i.e., the covariance depends only on the distance between two points, then the covariance matrix will be constant along the diagonals, a structure known as a Toeplitz matrix. If the timeseries wraps around and is isotropic, the structure is called a circulant matrix. Real life processes are seldom isotropic due to heteroskedasticity or edge effects. Heteroskedasticity, or unequal variance, will be visible along the diagonal of the covariance matrix. Likewise, edge effects can be seen at the corners of the diagonal of the covariance matrix. Many real-life timeseries in neuroscience are heteroskedastic or have edge effects, for instance, because the signal of interest is in the center of the timeseries. We will return to each of these special cases and examine them in depth.

A final benefit to our focus on the covariance matrix is that we do not have to assume any specific model was the generator for our data. For illustration and formal proofs, we must choose covariance matrices that correspond to specific statistical processes. However, in general, these results are not confined to just one process. For example, we do not need to assume that a processes is a Gaussian process. While a Gaussian process is one such process that would allow us to specify the covariance matrix, any statistical process with a given covariance matrix will lead to the results we present here. Therefore, our result are general across a wide range of statistical processes.

We will refer to *D*^*T*^ *D* as the covariance matrix of the mean-subtracted dataset *D*. Standard conventions define the covariance matrix as *DD*^*T*^, which assumes a column-vector layout of *D*. Since PCA uses the row-vector covariance matrix, we will mirror this convention.

#### 1.3 Singular value decomposition

Singular Value Decomposition (SVD) is a close cousin of PCA. The two are identical with one exception: the mean is not subtracted in SVD as it is in PCA, making PCA a special case of SVD. This relaxed assumption of SVD means that, rather than operating on the covariance matrix as in PCA, SVD operates on the matrix of non-central second moments of the columns of *D*, namely *D*^*T*^ *D* where *D* has non-zero mean. In practice, this usually means that the first singular vector (SV) of SVD resembles the mean, and the higher SVs are similar to the PCs. Most of our analyses will focus on SVD due to its convenient mathematical properties.

### 2 Smoothness-driven phantom oscillations

We consider two perspectives into smoothness-driven phantom oscillations. For a qualitative approach which generalizes into multiple dimensions, we show the equivalence between phantom oscillations and oscillatory modes through differential equations. For a more rigorous approach in the one-dimensional case, we compute the PCs using the Karhunen-Loève transform.

#### 2.1 Differential equations perspective

One way to conceive of smoothness in PCA is through the use of differential equations. This approach is attractive because it is visually-driven, conceptually easy to understand, and easily generalizes to multiple dimensions, such as the spatial cases. However, we use this approach here only to develop a qualitative sense of how phantom oscillations arise from smooth data, since it is more difficult to find exact solutions for the eigenvectors.

We begin by considering a one-dimensional continuum with band-limited isotropic smoothness. So, the covariance matrix can be parameterized by a function of a single variable, *f* (*d*), where *d* is the distance between two points. For convenience, we will assume that *f* is zero for distances greater than some distance *d*_*max*_. In this case, the covariance matrix will be a band matrix, a special type of Toeplitz matrix which has constant values along the central diagonals and zeros elsewhere. Let’s consider the case where the size of the matrix is large in comparison to *d*_*max*_.

Using this information, we can set up a relationship for all of the terms of an eigenvector *v* of length *N* which are more than *d*_*max*_ distance from the edge, with *N » d*_*max*_. For some eigenvector *v* with eigenvalue *λ*, we have,

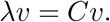

So then, for a non-edge point of the eigenvector *v*[*k*], such that *k* − *d*_*max*_ ≥ 0 and *k* + *d*_*max*_ *< N*, we can write it in a more illustrative form.

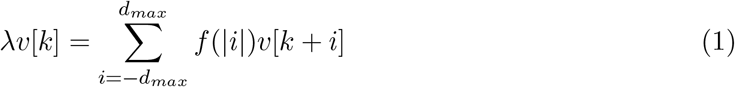

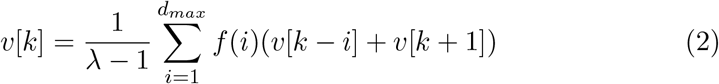

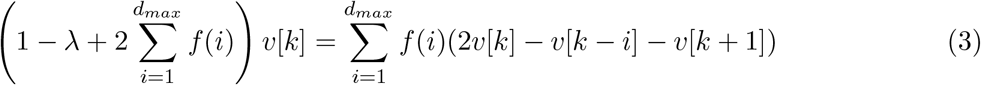

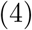

For any value of *i*, the term (2*v*[*k*] − *v*[*k* − *i*] − *v*[*k* + 1]) is the first-order finite central difference approximation of the second derivative. Conceptually, this is because the second derivative quantifies curvature, or how different a value is from the mean of its neighbors. Therefore, the weighted sum of these on the right hand side of the equation is also an approximation of the second derivative. So, if we write the continuous differential equation corresponding to this difference equation, using the continuous function *v*(*t*) and defining the constant 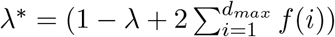, we have

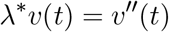

This is exactly the equation for a harmonic oscillator. Therefore, *v*(*t*) is satisfied by linear combinations of sines and cosines.

However, before interpreting *v*(*t*) directly as an eigenfunction, or a continuous equivalent of the eigenvector *v*, we must recall our initial assumption in deriving this equation, namely, that we are looking at points more than a distance *d*_*max*_ from the edges of the vectors. Consider the case exactly on the boundary where *k* = 0. Then, if our value of *λ*^*∗*^ remains the same, our derivation above becomes

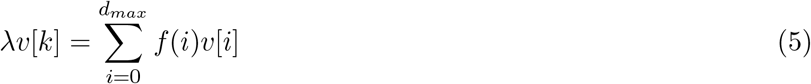

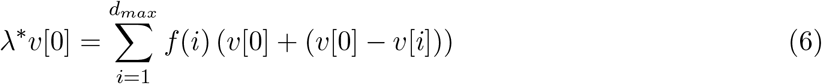

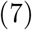

The term *v*[0] − *v*[*i*] is the (negative) first order forward finite difference approximation of the first derivative with a step size *i*. So, in the language of differential equations, the value at the edge of the eigenvector satisfies

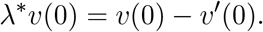

This derivation specifies the boundary conditions for the differential equation given above. The equation is a so-called “Robin boundary condition”, the combination of the more popular “Neumann” (first derivative equal to zero) and “Dirichlet” (fixed value) boundary conditions. While the Neumann and Dirichlet conditions specifiy that the boundary should be at the peak or zero-crossing, respectively, of the sine wave, the Robin condition specifies a value intermediate to the two. This means that subsequent frequencies may not fall on integer or quarter-integer intervals, as we would expect from the discrete cosine transform or the terms of a Fourier series. We will return to this idea later through other approaches.

We note that one special case of the above is the case where there are no boundaries. This occurs when the edges of the eigenvector have the same correlation as neighboring points, and is represented by a circulant matrix. In this case, we do not have to worry about the boundary conditions. Hence, every point is in interior point, and the eigenvectors are pure sines and cosines. We discuss this in more detail in the section “Wrap-around case”.

Lastly, while we considered only a single dimension, one additional benefit to the differential equations perspective is that it can be equivalently applied to the multidimensional case. It can also be applied to cases with complicated topologies, such as geodesic measurements on a brain surface, or positions on a complicated manifold described by a graph. In higher dimensions, the value of the function at a center point is proportional to a sum of estimates of the Laplacian, or the sum of the partial derivatives along each orthogonal dimension. (The above derivation can be applied directly for any geometry which exhibits 180^*°*^ rotational symmetry across all axes, which is true of most latices.) The solution to the continuous version of this equation are known as the Laplacian eigenfunctions, the harmonics, or the modes of vibration. This results in oscillatory-like or wave-like patterns in multi-dimensional space.

In summary, this analysis demonstrates that for smooth, isotropic, band-limited timeseries, the eigenvectors of the covariance matrix resemble sines and cosines away from the boundaries. However, they may have complicated boundary conditions which make it difficult to compute the eigenvectors analytically. The use of numerical tools can give the eigenvectors exactly, but cannot be used to prove their oscillatory behavior. For this, we will invoke the a generalized version of PCA, the Karhunen-Loève transform.

#### 2.2 Karhunen-Loève transform

The generalized, continuous version of PCA, the Karhunen-Loève transform (KLT), will form the basis for most of our proofs below. The KLT aims to find a sequence of functions on the interval [*t*_*min*_, *t*_*max*_] which maximally decorrelates a set of random processes on the same interval. These random processes may be discrete, but in the case we are most interested here, they will be continuous. While PCA operates on a specific discrete sample, KLT operates on a statistical process, where no such samples have been drawn. In this way, it is analogous to the PCs for an infinitely large sample with infinitely fine precision. PCA itself is a special case of the KLT for processes defined over discrete time which takes discrete values.

Just as PCA operates on the covariance matrix, KLT operates on the continuous analogue of a covariance matrix, the covariance kernel. As discussed above, a covariance kernel describes the expected value of the covariance of a sample drawn from any two given points in time. Likewise, the eigenfunctions of a covariance kernel are analogous to the eigenvectors of a covariance matrix. For a covariance kernel *C*(*t*_1_, *t*_2_), the *k*’th eigenfunction is function *v*_*k*_ : [*t*_*min*_, *t*_*max*_] → ℝ such that

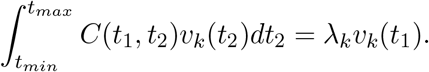

Note that this definition mirrors the multiplication of a covariance matrix *C* by an eigenvector *v*_*k*_[*i*], namely,

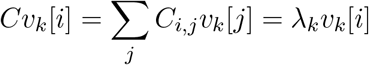

KLT states that the eigenfunctions *v*_*k*_(*t*) form an orthogonal basis of the Lebesgue function space *L*_2_[*t*_*min*_, *t*_*max*_] and that, for a stochastic process *X*(*t*), we have

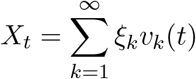

where *ξ*_*k*_ is a (not necessarily normally-distributed) random variable with mean 0 and variance *λ*_*k*_. Likewise, this is analogous to PCA, whereby a discrete timeseries *x*[*t*] can be written

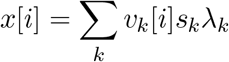

for the orthonormal vector of scores *s*.

While we can use fast linear algebra methods to compute exactly the eigenvectors of the co-variance matrix, finding a closed form expression for the eigenfunctions of a covariance kernel can be challenging. We review below some specific cases of smooth data for which the KLT has been solved analytically. In each of these cases, sinusoidal oscillations are produced.

#### 2.3 Specific solutions using the Karhunen-Loève transform

Using the KLT, we can solve for the eigenfunctions of several specific forms of the covariance kernel. These correspond to the eigenvectors of the corresponding covariance matrix, and hence, the principal component loadings of timeseries derived from an infinite number of samples of a given generative process. The principal components of these examples are shown in Figure S3 compared to those for white noise. We will not show the KLT derivations here, but will provide references.

##### Brownian motion

Brownian motion, diffusion, or a Gaussian random walk all produce similar covariance matrices. We assume that this process always starts at a given point, and runs from *t* = 0 until *t* = *T*. Note that this means Brownian motion is not isotropic. The covariance kernel is

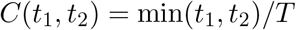

and the *k*’th eigenfunction has been found [73] to be

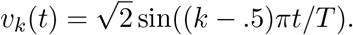

Likewise, a Brownian bridge is Brownian motion adjusted such that, in addition to the initial value being equal to zero, the value at time *t* = *T* is equal to zero. The covariance kernel is given by

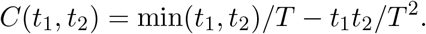

and the *k*’th eigenfunction has been found [73] to be

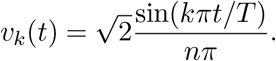

A similar construction applies to mean-centered Brownian motion [19].

Therefore, the PCs of Brownian motion and its variants have a sinusoidal form.

##### AR(1) and Ornstein-Uhlenbeck processes

A first-order autoregressive (AR(1)) process is one which each value is a weighted sum of the previous value and a random sample from a Gaussian distribution. It is generated with the formula

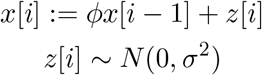

The covariance kernel for an AR(1) process, given sufficient time to make the initial conditions irrelevant, is

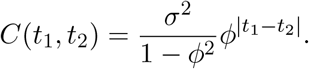

An Ornstein-Uhlenbeck process is a continuous version of an AR(1) process, and is given by the stochastic differential equation

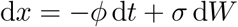

where *W* is a Wiener process. Ornstein-Uhlenbeck processes are closely related to dynamical systems perspectives of neural dynamics. The covariance kernel for the Ornstein-Uhlenbeck process, after enough time to avoid the effect of initial conditions, is

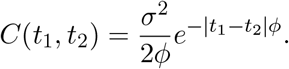

which can be rewritten in a form which looks more like that of the AR(1) process of a con, namely,

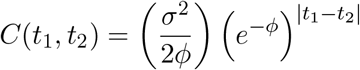

The eigenfunctions for *C* have an explicit solution [23, 32, 33, 73]. The *k*’th eigenfunction, defined on the interval [−*t*_*max*_, *t*_*max*_], for even *k* is

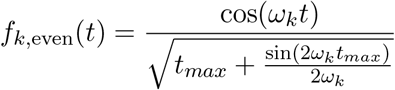

and for odd *k* is

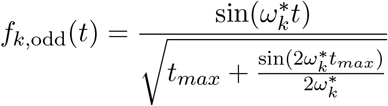

where *ω*_*k*_ is the *k*’th solution to the equation

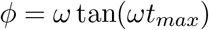

and 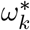 is the *k*’th solution to the equation

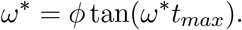

Note that these equations only depend on *t* within the sinusoids in the numerator. So, these eigenvectors are simply normalized sinusoids.

A similar derivation has been performed for an Ornstein-Uhlenbeck bridge, similar in concept to the Brownian bridge [14], and has also been solved in multiple dimensions [32, 33].

##### MA(1) or triangular kernel

A first-order moving average (MA(1)) process denotes a Gaussian random noise process smoothed with sliding window of length 2. It is generated by the process

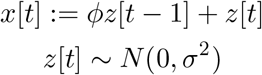

The covariance kernel is

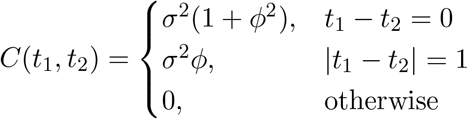

This is a special case which can be computed without interpreting the MA(1) process as a continuous function or invoking the Karhunen-Loève formalism. This forms a symmetric tridiagonal Toeplitz matrix, with (1 + *ϕ*^2^)*σ*^2^ on the diagonals and *ϕσ*^2^ on the off-diagonals. Using a recursive formula, it can be shown [45] that the *k*’th eigenvector of length *N* is

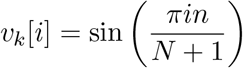

where *i* and *k* range from 1 to *N*. Note that the eigenvectors do not depend on the value of *ϕ* or *σ*. Also note that these functions are equivalent to the discrete sine transform [6]. So, all MA(1) processes, regardless of their parameters, will have the same eigenvectors.

While there is no single generalization of the MA(1) model to a continuous function, one option is a triangular covariance kernel. For some band-limiting parameter *T*, this has the form

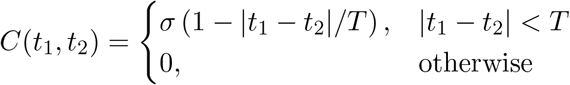

On the interval [0, *t*_*max*_], the *k*’th eigenfunction can be shown [23, 36] to be for even *k*

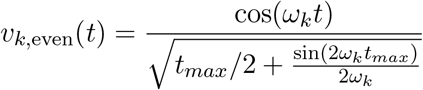

and for odd *k*

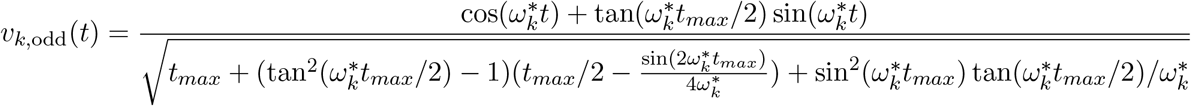

where *ω*_*k*_ is the solution to the equation

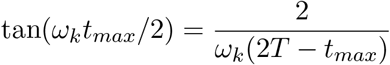

and

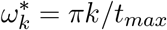

Notice that the only part of these equations which depend on *t* is the numerator, which is a single sinusoid in the even case and a sum of sinusoids with the same frequency in the odd case, allowing them to be written in the form *c*_0_ + *c*_1_ cos(*ω*_*k*_*t*) or 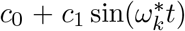 for the appropriate constants *c*_0_ and *c*_1_. Therefore, they are simply a shifted and rescaled sinusoid. Also notice that the odd solutions are fully independent of the band-limiting parameter *T*, and the even solutions depend on it only to set the underlying scale of the oscillatory frequency. Therefore, the width of the triangle kernel has minimal influence on the eigenvectors.

In practice, a very large sample size relative to the number of timepoints is necessary to estimate the covariance matrix of an MA(1) model with sufficient accuracy to produce phantom oscillations. Only higher-order MA models, triangle covariance kernels with a large bandwidth, or a small number of measured time points result in MA models having reliable phantom oscillations with reasonable sample sizes.

#### 2.4 More general results

While additional special cases have been explicitly derived using KLT [2, 23, 36, 52, 61, 73] or without KLT [17], it is nearly impossible to find a closed-form expression for the eigenvectors for all possible forms of a covariance matrix.

Prior work has permitted the approximation of the eigenvectors for large class of Toeplitz matrices [6], incorporating a broad class of covariance matrices for smooth data. A general symmetric Toeplitz matrix can be written as

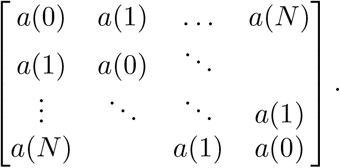

If the function *a* satisfies a “simple loop” smoothness constraint, then the eigenvectors of this matrix can be approximated as the sum of boundary conditions, a central component, and an error term. Roughly, the “simple loop” constraint requires that the inverse Fourier transform of the real-valued function *a*^*∗*^(*t*) = *a*(|*t*|) has a single maximum and a strictly monotonic decay towards zero on the unit circle. Then, the central portion of the eigenvector can be approximated by a sine function. Details of the proof are found in Bogoya et al. [6].

Lastly, in many cases, the oscillations of the eigenvectors may resemble functions other than sines or cosines, such as the Legendre polynomials [54]. While it is difficult to approximate these results analytically, several results specify the number of times the eigenfunction must cross zero. For instance, a proof in spectral graph theory using Fiedler’s nodal domains is given in Spielman [68].

#### 2.5 Wrap-around case

Here, we present a simple proof for the specific case of homogeneous spatial autocorrelation with even spacing in which the edges are connected (a torus). Suppose we have a discrete lattice of positions 1, …, *N* such that the covariance depends only on the distance between them with wrapping at the edges, i.e., for some function *f* (*d*), we have cov(*t*_*i*_, *t*_*j*_) = *f* (min(|*i* − *j*|, *N* − |*i* − *j*|)). Then, the covariance matrix *C* has the form of a symmetric circulant matrix, namely,

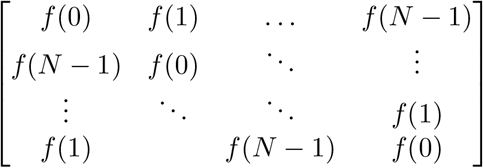

A well-known property of circulant matrices is that the *k*’th eigenvector is the *k*’th component of the discrete Fourier transform, namely,

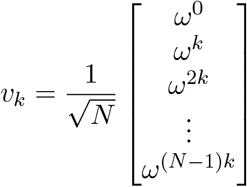

where *ω* = *e*^2*πj/N*^ is the *N* ‘th root of unity and 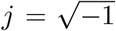. Furthermore, the corresponding *k*’th eigenvalue is

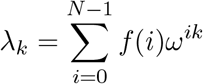

Since our circulant matrices are covariance matrices, they must be symmetric. If we consider the case of odd *N*, we have the relationship

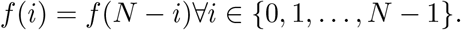

Using this, we can rewrite the terms for the eigenvalues as

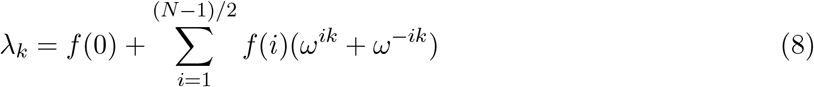

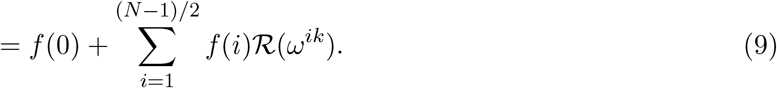

Notice that the eigenvalues are real, since the terms of *f* are real and *ω* always occurs in pairs of complex conjugates. Additionally, we see that the *k*’th eigenvalue is equal to the (*N* − *k*)’th eigenvalue for *k >* 0. This means that this system has repeated eigenvalues. An important property of repeated eigenvalues is that any linear combination of the associated eigenvectors is also an eigenvector.

This means we can write

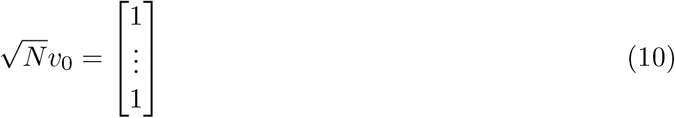

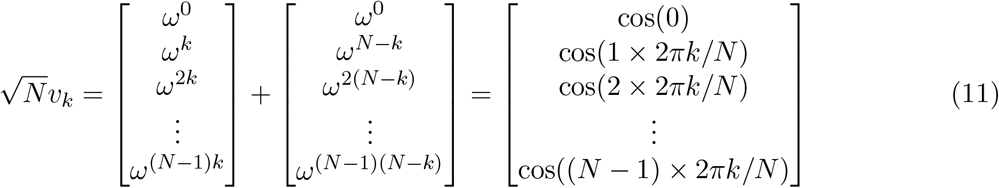

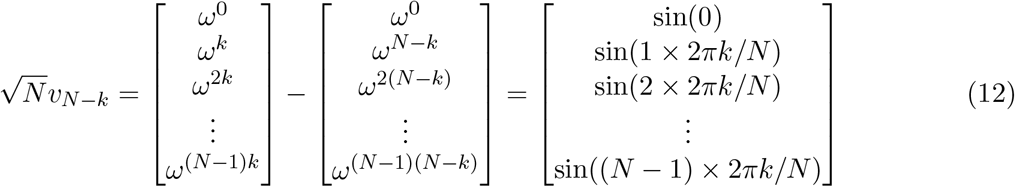

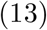

Therefore, the eigenvectors, and hence the loadings of the principal components, are pure sines and cosines.

For even *N*, the derivation is similar, except we have one additional eigenvector

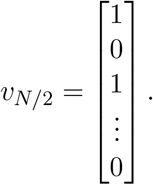

In this derivation, we used the well-known fact that the eigenvectors of a circulant matrix are given by the Fourier series. To conceptually understand the reasoning of why this is the case, we return again to the differential equations analysis of the Toeplitz matrix. We showed previously that, for a non-edge point *k* of an eigenvector *v* of length *N*, such that *k* − *d*_*max*_ ≥ 0 and *k* + *d*_*max*_ *< N*, then the eigenvector satisfies the relationship 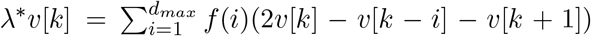. However, in the circulant matrix, this relationship also holds when the arithmetic is performed mod *N*. Therefore, every point in the eigenvector must satisfy this relationship. Unlike the case above, this means that there are no boundary points which satisfy a different equation. Thus, the continuous eigenfunction *v*(*t*) satisfies the relationship *λ*^*∗*^*v*(*t*) = *v*″ (*t*) across its entire domain. As a result, any solution to this differential equation which is cyclic over the interval [0, *N*] is an eigenfunction. The terms of the Fourier series are linearly independent and are the only real functions which satisfy this relationship. Therefore, sines and cosines are the eigenfunctions of a circulant matrix, regardless of the specific form of the matrix.

#### 2.6 Proof ideas that don’t work

There are at least two ways of approaching a proof about oscillatory eigenvectors which seem reasonable at first, but do not hold up to more rigorous mathematical analysis. First, if the covariance matrix is not a circulant matrix, it is not possible to prove that smoothness causes phantom oscillations by showing convergence to a Fourier series. Conceptually, this is an alluring approach: Fourier analysis is excellent at detecting oscillations. Furthermore, smoothness can be seen as a linear low-pass filter, represented by multiplication in the Fourier domain. So, it seems plausible that the lowest frequency terms of the Fourier series could serve as eigenfunctions.

While convergence to a Fourier series sounds reasonable, it does not work in practice because phantom oscillations do not match exactly the terms of the Fourier series. For a concrete example of the failure of Fourier analysis, we can see in Figure S3 that the phantom oscillations rarely occur in exact sine and cosine harmonics. Importantly, phantom oscillations my not have integer frequencies. A simple representation in Fourier space assumes that there are an integer number of cycles per vector. In the case of AR(1) or Ornstein-Uhlenbeck processes, as well as triangular kernels, we must solve a non-linear equation to find the number of cycles per vector, and in practice, these solutions are unlikely to be an integer. This causes the phantom oscillations to be split across multiple terms of the Fourier series, and so, the variance explained will also be split between these terms. Therefore, a Fourier transform is in general unable to find the same oscillations as those produced by PCA.

Second, it is not possible to prove that smoothness causes phantom oscillations by showing that a series of Toeplitz matrices converge to a circulant matrix in some infinite limit. Many of the covariance matrix forms we presented in Figure S3 are Toeplitz matrices. As the size of a banded Toeplitz matrix increases, it in some sense becomes more similar to a circulant matrix. Since we showed above that circulant matrices have eigenvectors equivalent to the Fourier transform of the circulant matrix coefficients, it is tempting to construct a series of Toeplitz matrices that converges to a circulant matrix in some limit. This strategy is effective for determining properties about the eigenvalues of Toeplitz matrices [26, 76], but not the eigenvectors. Gray [26] explicitly drew attention to the fact that their method applied only to eigenvalues, yet this confusion when citing Gray [26] remains a common mistake in the literature.

There are several conceptual issues which prevent the eigenvectors of a Toeplitz matrix from being approximated by the eigenvectors of a circulant matrix. For example, a circulant matrix ensures that the first element of the eigenvector *v* is related to the last element of *v*. But Toeplitz matrices in general do not do this. No reasonable definition of convergence can change the qualitative property of whether the first element of the eigenvector is related to the last element. In practice, this difference is manifest in the number of eigenvectors at each frequency. Circulant matrices contain two eigenvectors of each frequency, offset by 90^*°*^, which correspond to the real and imaginary parts of the Fourier transform. The linear combination of these allows any phase of sinusoid to be represented. In the differential equations view, circulant matrices have this property because the last term is related to the first term in the same way that all other neighboring terms are related, meaning there are essentially no boundaries or constraint on the phase of the oscillation. By contrast, the eigenvectors of Toeplitz matrices in general contain only one sinusoid at each frequency. The boundary conditions at the edges of the Toeplitz matrix constrain the phase of the oscillation, so they require only one sinusoidal eigenvector at each frequency. This discrepancy was the initial motivation for the discrete cosine transform [1, 27], a technique which is related to but distinct from the discrete Fourier transform.

### 3 Shift-driven phantom oscillations

In addition to generic smooth functions described above, we may also get oscillations in a case with a highly structured covariance matrix where a single signal is randomly shifted in time.

#### 3.1 Intuition

To gather an intuition, imagine we have a curve with one independent variable x and one dependent variable y, and a second identical curve shifted in the x direction by some small quantity *E*. Because PCA requires us to find the variance at each x position, we would like to find the distance between these curves in the y direction at each x position. Despite the fact that x is shifted by a constant amount *E*, the distance in y differs greatly at different points in the curve. Specifically, the x positions which have the largest slope are the ones for which y changes the most. This is because the definition of the slope of a function *f* (*t*) is “rise over run”, which in this case means (*f* (*t* + *ϵ*) − *f* (*t*))*/ ϵ*. Notice that as *E* gets smaller, this is the definition of the first derivative of the function *f* ! Also notice that if we shift the curve in the opposite direction, the sign of the derivative will flip. Therefore, if *E* is extremely small, we have a single function (namely, the derivative of *f*) which will tell us how far apart the original curve is from the shifted curve! If there were only two curves in the population, we would be able to explain nearly all of the variance with just the derivative.

Our intuition above can be extended to more realistic situations. In practice, *ϵ* will not be one single shift, but rather, a distribution of small but distinct shifts. But, the larger *E* is, the larger the difference between the curves will be. So, *ϵ* and the derivative jointly parameterize the magnitude of the difference between the curves. We can see this with the definition of the derivative: if the derivative is approximately constant for most *ϵ*, then due to *ϵ* being in the denominator, the difference between *f* (*t* + *ϵ*) and *f* (*t*) must be larger for larger *ϵ*. Slightly larger *ϵ* means that the derivative isn’t a perfect representation of the change anymore, because the derivative itself is changing and we need to take this into account. But consequently, we can perform this same analysis on the derivative of *f*, and find that the derivative changes the most when the second derivative of *f* is large. Thus, higher order derivatives are able to explain the difference between the curves.

#### 3.2 Formal analysis

We can more formally prove our intuition about the derivative. We first prove that to a first order approximation, a timeseries which is randomly shifted will have a first principal component equal to its derivative. Furthermore, if the shifts are normally distributed, in the second order approximation, the first two principal components will be equal to the first two derivatives. This proof is inspired by the proof given in Supplement 2 of [39].

Suppose we have many timeseries which represent the same signal *f* (*t*), but at slightly different time lags. Here, we will assume the timeseries are continuous functions. For simplicity, we assume that all timeseries are normalized. Without loss of generality, we can say |*f* (*t*)|_2_ = 1. Let the *i*’th timeseries *x*_*i*_(*t*) = *f* (*t* + *ϵ*_*i*_), where *ϵ*_*i*_ is a small positive or negative number. Thus, at a given time *t*, the vector-valued function *x*(*t*) = [*f* (*t* + *ϵ*_1_), …, *f* (*t* + *ϵ*_*n*_)]. Therefore, the covariance between two different timepoints is

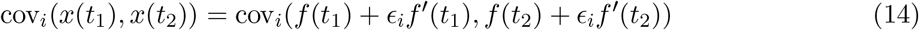

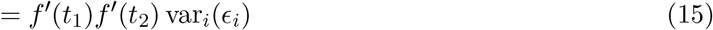

When measured across several timepoints *t*_1_, …, *t*_*n*_, we can form the vectors **f** = [*f* (*t*_1_), …, *f* (*t*_*n*_)], as well as the derivatives **f**′ = [*f*′ (*t*_1_), …, *f*′ (*t*_*n*_)] and so on.

The covariance matrix is thus

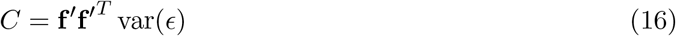

Since this is a rank-1 matrix, it is trivially diagonalizable, and hence, **f**′ is an eigenvector of the covariance matrix *C* with eigenvalue var(*ϵ*).

To see the impact of higher-order derivatives, we can extend this to higher moments in the Taylor series. For the first three terms of the series

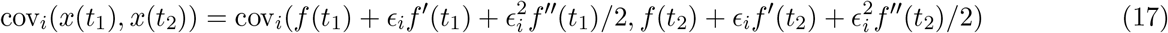

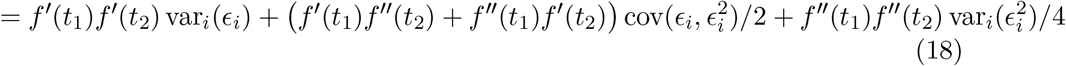

or equivalently

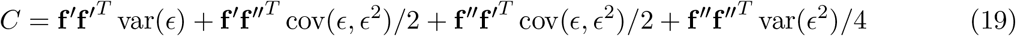

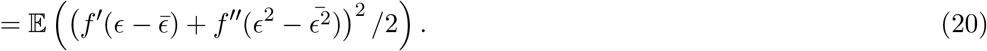

We see that the covariance matrix can be written as a sum of terms based on the higher derivatives.

If we make the further assumption that *ϵ* ∼ *N* (0, *σ*^2^), then we can simplify this expression further. For *k* ∈ ℕ_0_, the (2*k* + 1)’th moments of a normal distribution with mean zero are equal to zero [74]. Thus, for large *N*, the term cov(*ϵ, ϵ*^2^) = 𝔼 (*ϵ*^3^) − 𝔼 (*ϵ*) 𝔼 (*ϵ*^2^) = 0. Therefore, we can write,

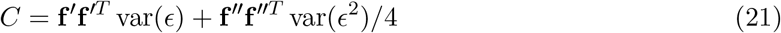

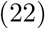

This means that the covariance matrix is diagonalizable with the following:

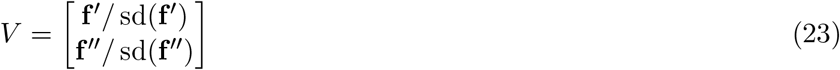

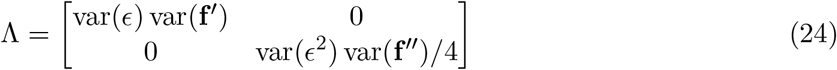

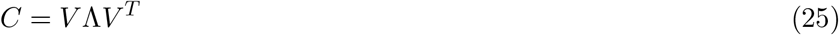

Thus, **f**′ and **f**″ are eigenvectors of *C*, and var(*ϵ*) var(**f**′) and var(*ϵ*^2^) var(**f**″)*/*4 are eigenvalues.

Note that if we look at the impact of the third derivative by adding a fourth term to the Taylor series, the simple diagonalizable form no longer holds. The covariance matrix then becomes

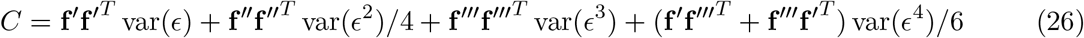

which permits a representation of

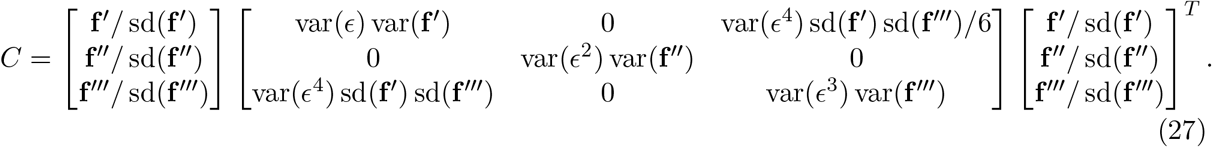

However, error at the magnitude of *O*(*ϵ*^4^) is unlikely to be biologically meaningful or scientifically interpretable.

Also note also that for larger time shifts, the Taylor expansion may not be a good approximation. A separate mathematical framework rooted in singular spectrum analysis (SSA) can analyze the covariance matrices of lagged timeseries [72].

### 4 PCA of the transpose

So far, we have focused on the case where the feature dimension is the continuous dimension. For instance, for the *m × n* data matrix *D*, this means that the rows consist of *m* observations, and the columns consist of *n* timepoints. However, it is also common to see the opposite case, where PCA is performed on the *n × m* matrix *D*^*T*^. If the observations dimension is continuous and the feature dimension is not, then the covariance matrix would show no obvious visible patterns. Nevertheless, phantom oscillations still occur when the matrix is transposed, but instead of appearing in the loadings, they appear in the scores.

If PCA performed on a matrix produces phantom oscillations in the loadings, then PCA performed on the transpose of the matrix will produce phantom oscillations in the scores. Suppose we have a data matrix *D*. Using singular value decomposition (SVD), or more specifically”full SVD”, we can rewrite this as

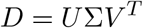

where *U* and *V* are orthonormal matrices of size *m × m* and *n × n* respectively, and σ is a *m × n* diagonal matrix padded with zeros. PCA is closely related to SVD: if the data matrix *D* has a mean of zero for each feature, then we can write the covariance matrix as *D*^*T*^ *D*. This means that we can write

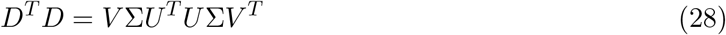

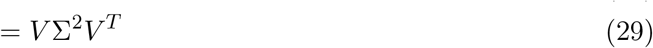

where Σ^2^ is and *n × n* diagonal matrix. Therefore, the PCs are given by the rows of *V* and the variance explained of each is given by Σ^2^. Furthermore, by substituting Equation 4, we see that the projection of *D* onto *V* is equal to *U* Σ. So, the PC scores are proportional to *U*.

If we perform PCA in the opposite direction, we can formulate it using SVD. Using the identities above from SVD, we can substitute and find

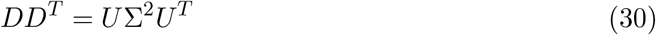

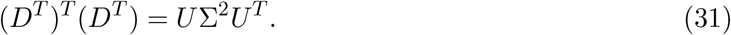

where Σ^2^ is an *m × m* diagonal matrix. (We use the same notation of Σ^2^ in each case because the values along the diagonal are identical, but one version is padded by zeros.) So now, we have a form for the SVD of *D*^*T*^ given the SVD of *D*. In this formulation, we can see that the oscillatory structure now occurs in the PC scores instead of the PCs themselves. In other words, these oscillations will still be present even if PCA is performed on its transpose.

Note that PCA is only exactly equal to its transpose (swapping loadings and scores) if has a mean of zero in both dimensions. PCA only requires the mean to be subtracted along the feature axis, not the observations axis, meaning *DD*^*T*^ is not a covariance matrix. Nevertheless, the eigenvalues of Σ^2^ are the same in both cases, and the first eigenvector is usually very similar to the mean [10].

Despite the lack of perfect equality due to mean subtraction, phantom oscillations in one dimension will lead to phantom oscillations in the other. [10] showed that the difference between the covariance matrix and the matrix of second moments is at most rank one. Therefore, the difference between covariance matrices for *D* and *D*^*T*^ is at most rank two, regardless of the size of *D*. However, as we showed above, phantom oscillations are present with steadily increasing frequency. Therefore, at most oscillatory components could be absorbed by differences in mean subtraction. For reasonably sized matrices, there will be more than two oscillatory components. Therefore, if phantom oscillations are present, they will be present regardless of any mean subtraction.

### 5 Artificial and experimental data

#### 5.1 Smoothness-driven phantom oscillations

The smooth artificial timeseries were generated as smoothed white noise. We generated 100,000 artificial timeseries of length 300 drawn from the standard normal distribution. We filtered with a Gaussian filter of standard deviation 4. We discarded all but the central 60 timepoints to avoid edge effects of the filtering, leaving 100,000 timeseries of length 60. We repeated this procedure to create an equally-sized dataset for cross-validation.

The resting state fMRI data were derived from the Cam-CAN project [64]. We used the resting state scans with the AAL parcellation, as provided by the Cam-CAN project. To remove transient high-frequency artifacts, we applied a low-pass Butterworth filter at half the Nyquist frequency. For each of the 646 subjects, we selected 60 parcels with replacement, and for each selection, we chose a random segment of 60 time points. Sampling rate (TR) was 1.97 s, for a total of 118.2 s of data. In total, this yielded 38760 timeseries. We randomly selected half of these, 19380, for our core dataset, and used the rest for cross-validation.

The spatially smooth data were generated by drawing from a multivariate normal distribution. Each timepoint was one sample from this distribution. The covariance matrix was determined as negative exponential of distance for each region or pixel. For the data in the geometry of a widefield recording, we drew 10000 independent samples of size 5916 (for 5916 pixels in the image) and for the geometry of cortex, 3000 independent samples of size 180 (for 180 pixels in the parcellation of Glasser et al. [24].

#### 5.2 Shift-driven phantom oscillations

The random shifts in timing data were generated based on the response of frontal eye field neurons to a delayed stimulus [67]. The base timeseries was taken from the freely-available data provided by Shinn et al. [67], using the mean activity from cells in Monkey Q with 70% coherence in the 400 ms presample condition, with data from 600 ms before and after stimulus onset with 10 ms time bins. The resulting timeseries was smoothed with a Savitzky-Golay filter of order 1 with a window length of 5. We drew 2000 time shift offsets from a uniform distribution from 0.1 to 0.4. The base timeseries was upsampled through linear interpolation according to each shift, and the segment from 0 to 700 ms was selected. This yielded 2000 timeseries of length 700. We repeated this sampling and interpolation process to create another 2000 timeseries to be used for cross-validation only.

The random shifts with non-identical signals were generated using noisy difference of gammas. We generated base data using

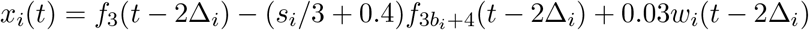

where Δ_*i*_ ∼ *U* [0, 1] is the shift in time, *b* ∼ *U* [0, 1] is the shape parameter, *s*_*i*_ ∼ *U* [0, 1] is the amplitude of the second gamma for timeseries *i, f*_*k*_(*t*) is the probability density function of the gamma distribution with shape parameter *k*, and *w*_*i*_(*t*) is 1*/f*^*α*^ noise with exponent 1.8. We chose this form arbitrarily because it visually approximates a noisy hemodynamic response function. We generated 2000 timeseries of length 10 s at 100 hz this way, and 2000 additional timeseries for cross-validation only.

The images were generated with a single static two-photon image of two neurons. The shifts for the image were computed by filtering a white noise timeseries of length 10000 with a Gaussian filter of standard deviation 5. Only a small segment of this timeseries is shown in (Figure 3g). Shifts were applied to the image, simulating movement of the imaging window, and images were flattened before computing PCA.

#### 5.3 Random dot motion task

Full details of the random dot motion experiment are described in Roitman and Shadlen [57]. Due to the limited number of neurons recorded in this dataset, analyses presented here pool data from both monkeys. For illustration purposes, in the schematic Figure 4b, we used neural activity averaged across all trials with an RT greater than 800 ms, with a bin size of 50 ms.

For our analysis of diffusion-like activity showing smoothness-driven phantom oscillations, we examined the period from 0 to 800 ms after the onset of the stimulus, using a bin size of 80 ms. We restricted to trials which had an RT greater than 800 ms. This gave a total of 1821 trials with a 10 time points per trial.

For our analysis of transient activity showing shift-driven phantom oscillations, we examined the period 500 ms before the saccade to 300 ms after the saccade, with a time bin of 50 ms. Because these transients generally occur when the target is in the receptive field, we only examined saccades to T-IN. To ensure a strong transient, we only considered trials with a motion coherence greater than 10%. Assuming each cell had a similar delay, we averaged the response of each neuron, for a total of 54 timeseries (one for each neuron) with 16 timepoints.

#### 5.4 Pedal task

Full details of the pedal experiment are described in Ames and Churchland [3]. We used data only from monkey E, starting from the top, pedaling in the forward direction with both hands. Activity was averaged for each neuron, and all sessions were pooled for a total of 626 neurons. While pedaling on each trial was performed at approximately the same speed, it was not exactly the same on each trial. Therefore, to find average firing rate for each neuron at each point in time, trials were aligned according to the pedal position, and this was rescaled across time such that each cycle took 500 ms. The first two cycles and the last cycle were discarded. The hand position example in Figure 5b uses the hand position of an example aligned trial.

## Supplemental figures

**Figure S1:**
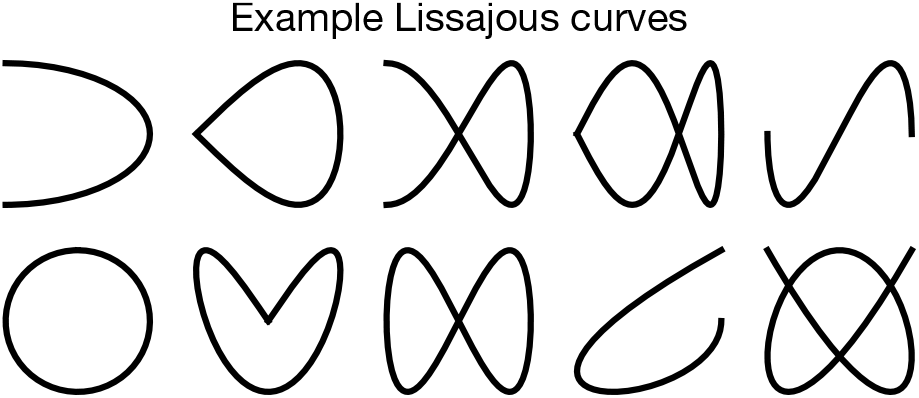
Simulated examples of neural trajectories. When PCs containing phantom oscillations are plotted against each other, they create Lissajous-like trajectory patterns common in neural state space. Shown here are several examples.

**Figure S2:**
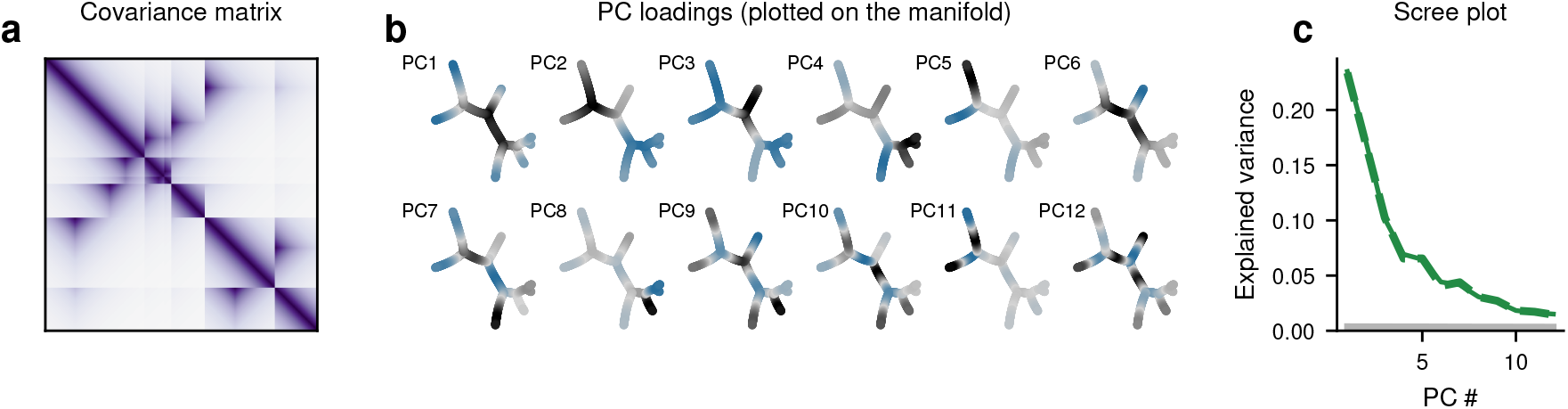
Phantom oscillations along a continuous manifold. Phantom oscillations may occur when continuity is along a branching manifold in multiple dimensions instead of across time. (a) Covariance matrix across observations, where covariance is a decreasing function of distance along the manifold’s graph. (b) The PC loadings. Since there is not one single continuous axis, loadings are displayed by color on a two-dimensional projection of the manifold. (c) A scree plot showing variance explained by each principal component.

**Figure S3:**
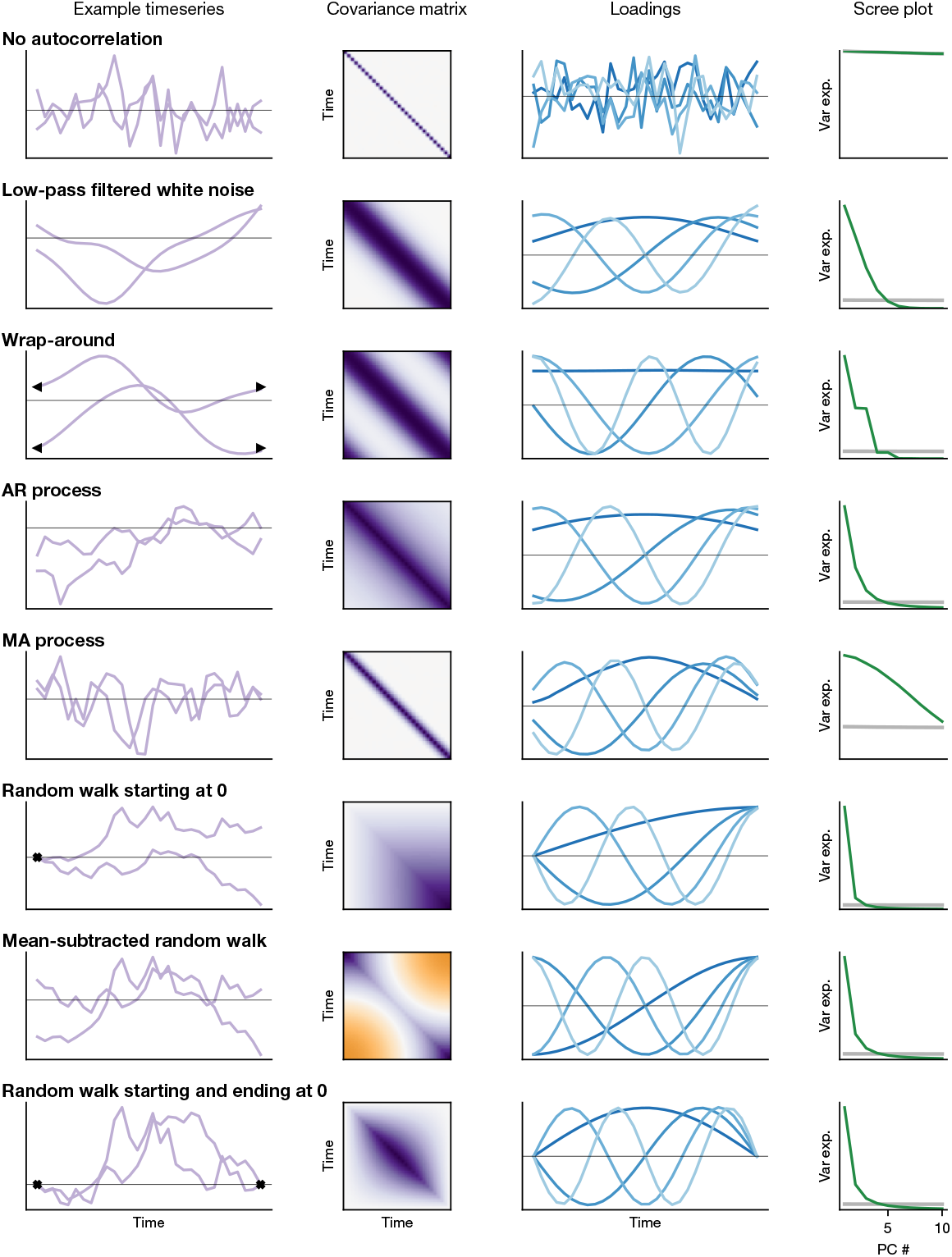
Different types of smoothness show different patterns of phantom oscillation. (left) For each process, two example timeseries are shown. Triangles indicate continuity between the two sides of the graph. X’s indicate the timeseries is anchored at this point in time. (left center) The covariance matrix for the process is shown, where purple indicates high covariance, orange indicates highly negative, and white indicates zero. (right center) The loadings of the first four principal components (PCs) are shown, where the center line indicates zero. The darkest shade of blue indicates PC1, and the lightest PC4. (right) The variance explained of the components for each example process. The grey line indicates the variance explained by the PCs of equally-sized white noise. For reference, the first process is uncorrelated white noise, showing no phantom oscillations. AR is a first-order autoregressive process, and MA is a moving average process.

**Figure S4:**
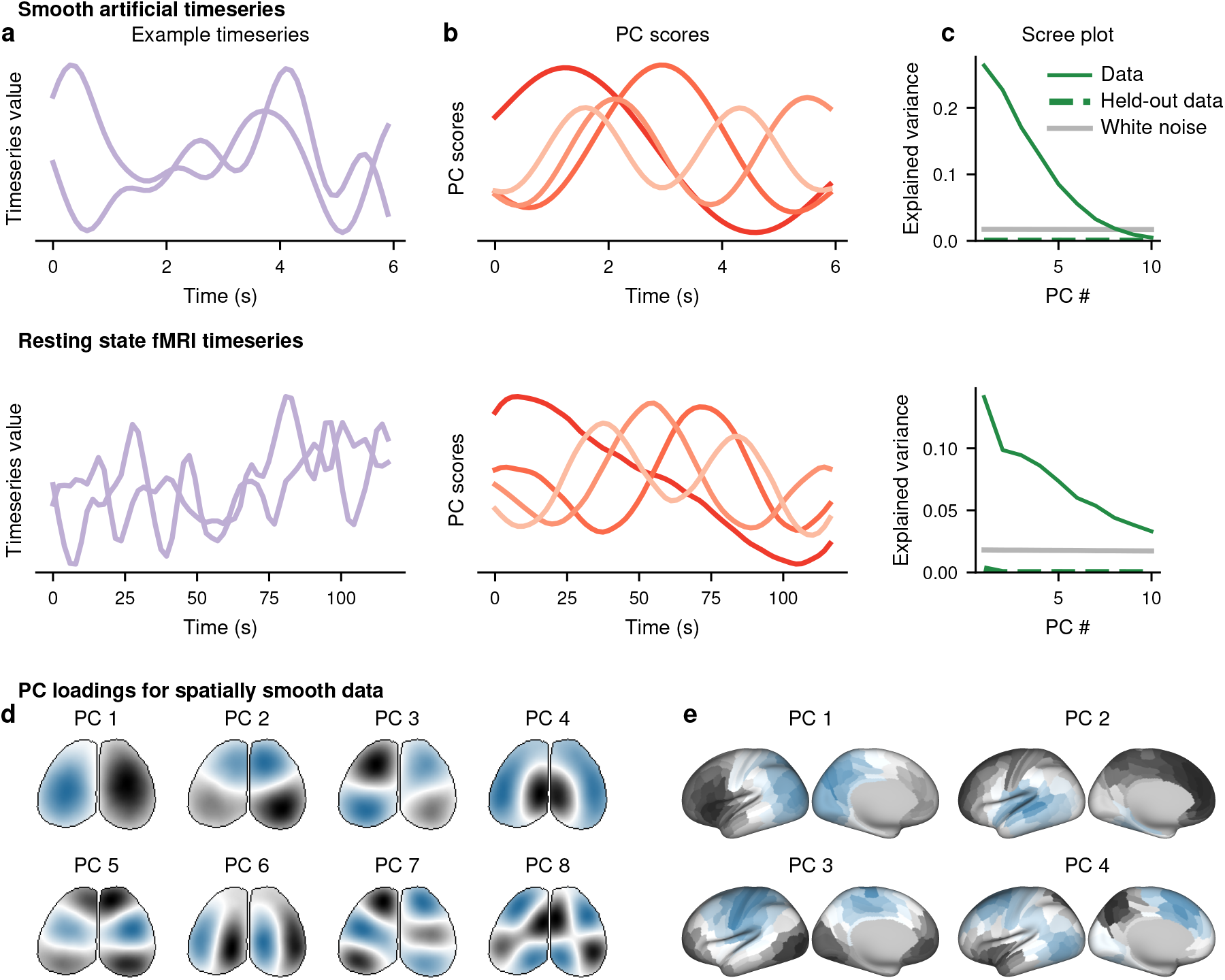
Smoothness causes phantom oscillations in PCA scores when time is in the observation dimension. Same format to Figure 2, except data matrix has been transposed, causing the smooth timeseries to be in the observation dimension and individual neural signals to be in the feature dimension. We do not show the loadings in a-c or the scores in Figure 2a-c because there is no apparent pattern. Held-out data explain little variance in c because the phantom oscillations occur in the scores rather than the loadings. A different cross-validation procedure, one which examined the similarity of the scores, would show cross-validation performance in c similar to Figure 2c.

**Figure S5:**
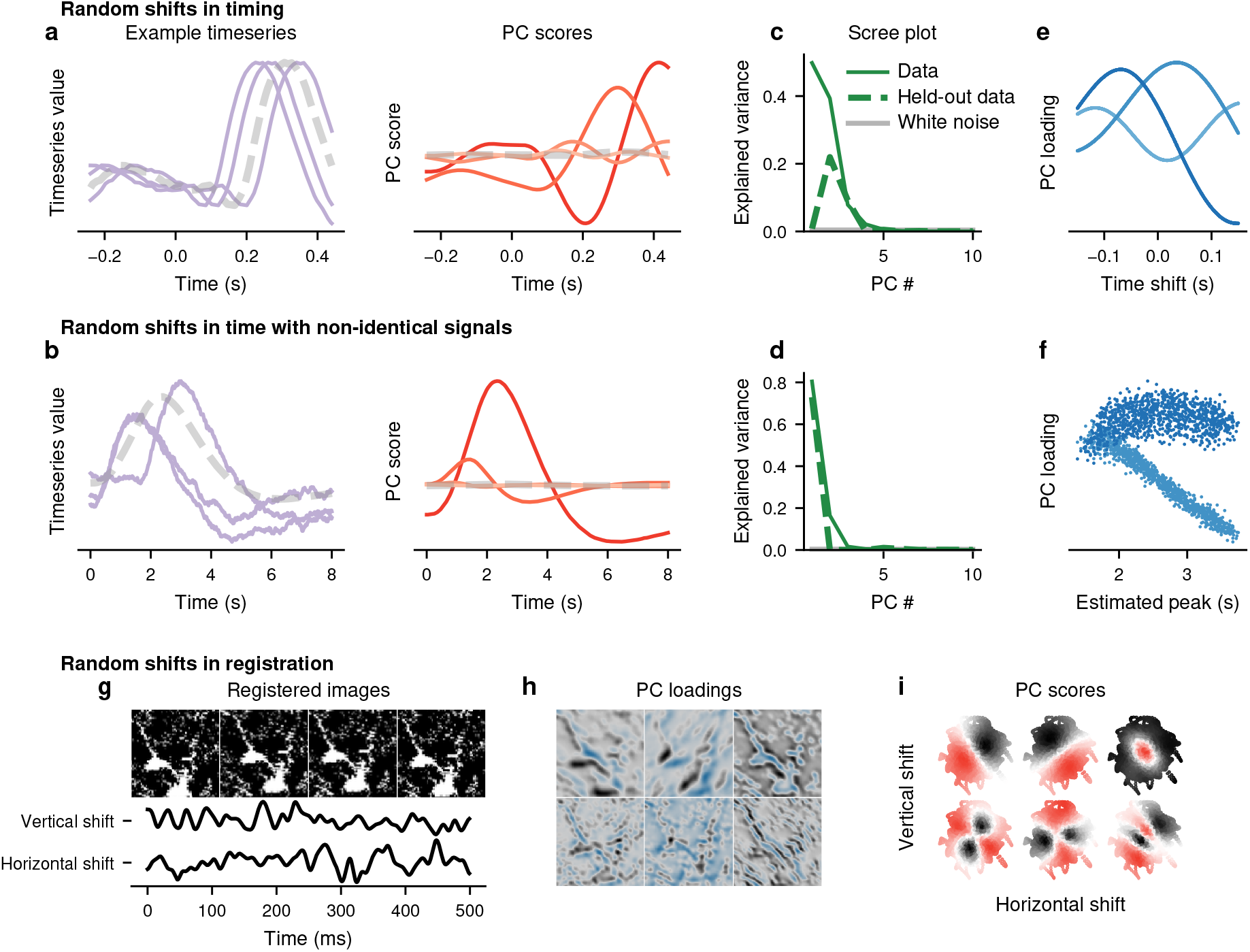
Time shifts cause phantom oscillations in PCA scores. Same format to Figure 3, except the data matrix has been transposed, causing the shifted timeseries to be in the observation dimension and individual neural signals to be in the feature dimension. Held-out data explain less variance in c-d because the phantom oscillations occur in the scores rather than the loadings. A different cross-validation procedure, one which examined the similarity of the scores, would show cross-validation performance in c-d similar to Figure 3c-d.

**Figure S6:**
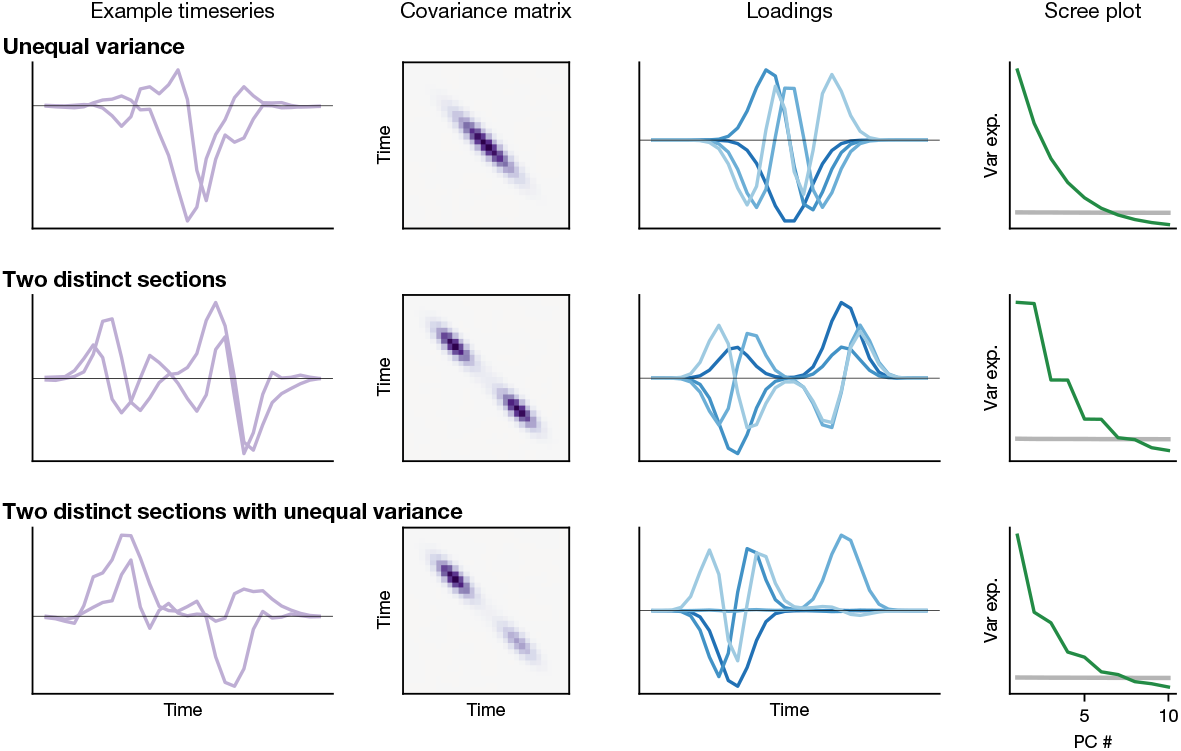
Simulated examples of different types of data. (left) For each process, two example timeseries are shown. (left center) The covariance matrix for the process, where purple indicates high covariance, orange indicates highly negative, and white indicates zero. (right center) The loadings of the first four principal components (PCs), where the center line indicates zero. The darkest shade of blue indicates PC1, and the lightest PC4. (right) The variance explained of the components for each example process. The grey line indicates the variance explained by the PCs of equally-sized white noise.

## Notes

### Competing Interest Statement

The authors have declared no competing interest.

